# NREM Oscillations Mediate Synaptic Proteome Remodelling to Support Synapse Stabilisation

**DOI:** 10.64898/2026.04.07.716883

**Authors:** Diego del Castillo-Berges, Ana Cazurro-Gutierrez, Oriana Zerpa-Rios, Amalia Peñuela, Daniel Arco-Alonso, Carla Viñola-Renart, Gisela Espriu-Aguado, Danique Zantinge, Thomas Vaissière, Camilo Rojas, Frank Koopmans, Remco V Klaassen, Benito Domínguez-Velasco, Manuel Alvarez-Dolado, Julie Seibt, Gavin Rumbaugh, August B Smit, Àlex Bayés

**Affiliations:** Molecular Physiology of the Synapse Laboratory, Institut de Recerca Sant Pau (IR Sant Pau), Barcelona, Spain; Universitat Autònoma de Barcelona (UAB), Bellaterra (Cerdanyola del Vallès), Spain; Institut Recerca Vall d’Hebrón, Barcelona, Spain; Institute of Biotechnology and Biomedicine (IBB) and Department of Cell Biology, Physiology and Immunology, Universitat Autònoma de Barcelona, Bellaterra (Cerdanyola del Vallès), Spain; Surrey Sleep Research Centre, School of Biosciences, Faculty of Health and Medical Sciences, University of Surrey, Guildford GU2 7XP, U.K; Department of Neuroscience, The Herbert Wertheim UF Scripps Institute for Biomedical Innovation & Technology, Jupiter, FL, USA; Department of Molecular and Cellular Neurobiology, CNCR, VU University and UMC Amsterdam, 1081 HV Amsterdam, the Netherlands; Andalusian Centre for Molecular Biology and Regenerative Medicine (CABIMER), CSIC-JA-US-UPO, Seville 41092, Spain

**Keywords:** Syngap1, sleep/wake cycles, NREM sleep, synapse, proteomics, brain oscillations, electroencephalography, spindles, slow-waves, mouse model, neurodevelopmental disorder

## Abstract

Synapses are known to remodel their proteome during sleep. However, the exact mechanisms driving this remodelling and its impact on synaptic function or cognition are not well understood. We combine 24-hour EEG recordings with time-resolved synaptic proteomics in a model of *SYNGAP1*-Related Disorders to reveal a mechanism by which slow-waves and spindles, two NREM sleep oscillations, mediate the remodelling of the synaptic proteome. Moreover, we uncover that this remodelling promotes synaptic stabilization, which could support sleep-dependent memory consolidation. In contrast, the increase of slow-waves and decrease of spindles found in *Syngap1*^+/-^ mice would activate molecular pathways involved in synaptic weakening instead of stabilization. This is consistent with the proposed roles of slow-waves and spindles in synaptic downscaling and potentiation, respectively. Here, we provide evidence on how NREM oscillations regulate the synaptic proteome and reveal a pathological mechanism that could be of relevance to all neurodevelopmental disorders coursing with sleep disturbances.

## INTRODUCTION

Sleep is an evolutionarily conserved process^1,2^ that serves essential biological functions. These include cognitive processes, such as learning or memory consolidation^3–6^, or cellular restoration events, such as metabolic waste clearance^7,8^. Consistently, sleep alterations lead to impairments in neural processing and behavior^9–11^. Furthermore, sleep disturbances are highly prevalent across neurological and psychiatric conditions, being particularly prominent in genetic neurodevelopmental disorders (NDDs)^12–15^. These observations highlight sleep as a critical component of brain health and a potential contributor to disease-related neural dysfunction.

Several studies have shown that sleep influences the proteomic composition of synapses^16–20^. Indeed, the synaptic proteome undergoes daily remodelling that are linked to sleep and wake cycles^18^. Importantly, these daily proteome dynamics are not caused by tissue-wide changes in gene expression or protein turn-over, instead, they are restricted to synapses^18^, and thus should be driven by synapse-specific mechanisms. This suggest that the marked differences in brain activity occurring between sleep and wakefulness lead to distinct patterns of synaptic stimulation, resulting in the remodeling of the synaptic proteome. Therefore, it is plausible that specific brain oscillations within a given state, such as slow-waves (SWs) and spindles in NREM sleep, are also able to influence synaptic composition in different manners. Nevertheless, the exact influence of different NREM oscillations onto the synaptic proteome remains unexplored.

It is well-known that both SWs and spindles play an important role in cognition, since their interaction facilitates the transfer of information from short- to long-term storage, contributing to memory consolidation during sleep^3,4,21^. SWs are oscillations in the 0.5-4 Hz range that emerge from alternating periods of firing and silence in cortical neurons^22^, while spindles are waxing and waning oscillations in the 9–16 Hz range that originate in thalamocortical circuits^21^. Both oscillations have been proposed to participate in synaptic plasticity. SWs would promote synaptic downscaling during sleep^23^, indeed, different patterns of stimulation delivered onto pyramidal neurons during up states produce synaptic weakening^24–26^. On the other hand, spindles closely correlate with calcium transients in cortical dendrites, suggesting they may facilitate synaptic potentiation^27–29^. Accordingly, a theoretical framework of sleep function has proposed that spindle-rich sleep supports synaptic and circuit potentiation, while SWs-rich sleep promotes homeostatic synaptic downscaling^30^.

Pathogenic variants in the *SYNGAP1* gene cause an autosomal dominant NDD involving intellectual disability and epilepsy^31–34^. Beyond these hallmark features, sleep disturbances are highly frequent in SYNGAP1-Related Disorders (SYNGAP1-RD), affecting about two-thirds of these individuals^32,33^. Importantly, among the sleep disturbances identified, we have recently shown that individuals with this condition present a marked decrease in spindle activity^33^. Alterations in this oscillatory activity have been reported in related NDDs, including Rett^35^, Angelman^36^, and rodent models of Fragile X^37^ or Dravet^38^. At the molecular level, the *SYNGAP1* gene encodes for a Ras/Rap GTPase-activating protein (SynGAP) highly enriched in synapses^39,40^, where it plays important functions in both Hebbian and homeostatic synaptic plasticity^41–45^.

Given the high prevalence of sleep disturbances in SYNGAP1-RD, the central role of the SynGAP protein in synaptic physiology, and the emerging evidence linking sleep to the regulation of the synaptic proteome, we asked whether *Syngap1* heterozygosity disturbs daily synaptic proteome dynamics. Here, we integrate 24-hour electroencephalographic (EEG) recordings with time-resolved synaptic proteomics in *Syngap1*^+/-^ mice to provide evidence linking slow-waves and spindles to synaptic proteome remodelling. We reveal a potential pathological mechanism in which disrupted brain oscillations would lead to dysregulated synaptic proteome dynamics. This could have important cognitive implications for NDDs and other neurological conditions with aberrant brain oscillations.

## RESULTS

### Sleep Architecture and Circadian Behaviour are Preserved in *Syngap1*^+/-^ Mice

We first performed 24h video recordings in wild-type (WT) and *Syngap1*^+/-^ (HET) mice to track their locomotor activity. These started at Zeitgeber Time (ZT) 0, that is, at the beginning of the light/rest period. As expected for nocturnal animals, both genotypes presented clearly enhanced locomotion during the dark/active period (**Figure 1a,b and S1a**), suggesting a well-preserved circadian behaviour. However, as previously reported^46,47^, *Syngap1*^+/-^ animals had a larger light to dark increase in activity (**Figure 1a,c**).

**Figure 1.**
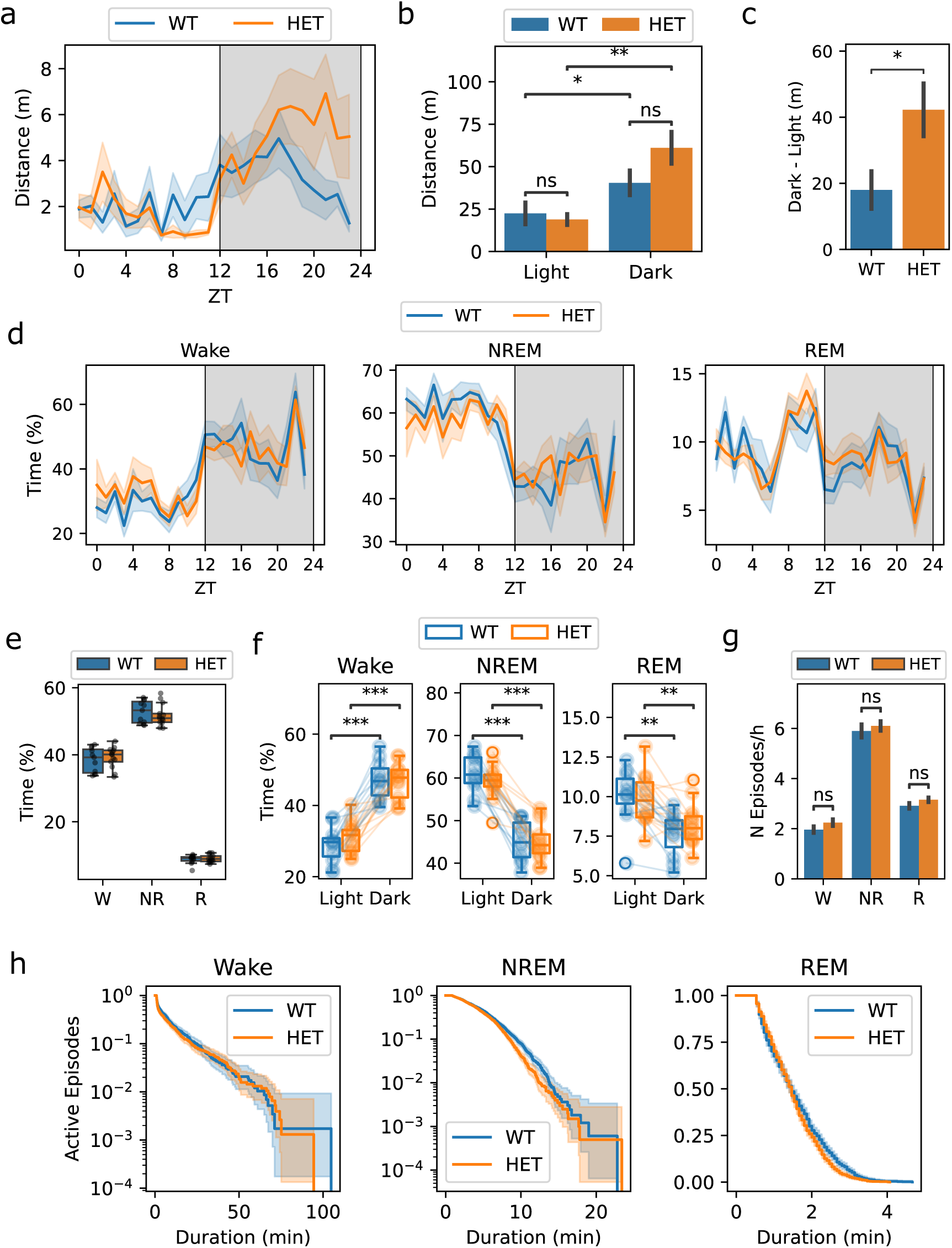
*Syngap1*^+/-^ mice show preserved circadian regulation of locomotor activity and normal sleep architecture. **(a)** Mean distance (m) travelled per hour over a 24-hour period, shown for wild-type (WT n=8, blue) and heterozygous (HET n=8, orange) mice. Shaded areas surrounding the means correspond to the standard error of the mean (SEM). Gray shaded area indicates dark period. Time expressed as Zeitgeber Time (ZT). **(b)** Mean distance travelled during light and dark periods for WT (blue) and HET (orange) animals. Data presented as mean ± SEM. A Mixed-ANOVA with between-subject variable ‘Genotype’ and within-subject ‘Circadian Period’ variable revealed a significant effect of the ’Circadian Period’ (Light vs Dark; p = 2.5e-5) and a significant Genotype x Circadian Period interaction (p = 0.027). Post-hoc analysis was performed using paired t-tests to compare light and dark period within genotypes (WT adj-p=0.047; HET adj-p =0.0044) and independent t-tests to compare between genotypes in each circadian period (Light adj.-p=0.21, Dark adj-p=0.62). Post-hoc contrasts were corrected using Holm correction. **(c)** Change in activity levels (Dark-Light) for WT and HET animals. Data presented as mean ± SEM. Statistics: independent t-test, p=0.027. **(d)** Hourly distribution of vigilance states across the 24-hour cycle for WT (n=11, blue) and HET (n=13, orange) mice: Wake (left), NREM (middle), and REM (right). Lines represent the mean and shaded areas indicate the SEM. Gray shaded area indicates dark period. **(e)** Percentage of total time spent in Wake (W), NREM (NR), and REM (R) sleep across 24 hours for WT (n=11, blue) and HET (n=13, orange) mice. No significantly differences found. Statistics: independent t-tests to compare Wake (p=0.59) and NREM (0.47) and Mann-Whitney to compare REM (p=0.91) **(f)** Percentage of time spent in each vigilance state (Wake, NREM, REM) during the light or dark periods for each genotype. Paired individual values are shown as dots connected by lines. Boxplots represent group distribution. Statistics: Mixed-ANOVA with ‘Genotype’ as between-subjects variable and ‘Circadian Period’ as within-subject variable for each state. ‘Circadian Period’ was significant in all state (W p=5.9e-11, NR p=1.13e-11, R p=1.2e-5). Post-hoc analysis within genotypes was performed using a paired t-tests with Holm correction (W: WT adj-p=7.93e-6, HET adj-p=7.93e-6; NR: WT adj-p=3.2e-6, HET adj-p=2.53e-6; R: WT adj-p=0.001, HET adj-p=0.008). **(g)** Number of episodes per hour and state: Wake (W), NREM (NR) and REM (R). Bars represent mean ± SEM across animals. Statistics, unpaired t-tests or Mann-Whitney for non-normal data. ns, non-significant. **(h)** Kaplan–Meier survival curves for Wake (left), NREM (middle), and REM (right) episodes, reflecting the proportion of bouts that remain active as a function of duration. Shaded areas represent 95% confidence intervals. No significant differences were found between genotypes: Cox-Regression Analysis, Wake: HR= 1.10, p=0.16; NREM: HR= 1.14, p=0.09; REM: HR= 1.15, p=0.09. Significance levels, *p < 0.05, **p < 0.01, ***p < 0.001, ****p < 0.0001.

To directly assess sleep-wake dynamics we conducted 24h EEG recordings (**Figure S1b,c**). First, we quantified the percentage of daytime spent awaken, in non-rapid eye movement (NREM) or REM sleep and found no differences between genotypes for any state (**Figure 1d,e**). Furthermore, both genotypes showed higher Wake amount during the dark/active period and higher NREM and REM amounts during the light/rest period (**Figure 1d,f**). Finally, we also studied sleep consolidation by looking into in the number and length of sleep and wake episodes (see Methods). We found no differences between genotypes in the number of episodes from each state (**Figure 1g**) or in the distribution of their lengths (**Figure 1h**). Overall, these findings indicate that our model of SYNGAP1-RD does not present substantial alterations in sleep architecture and circadian behaviour.

### *Syngap1* Heterozygosity Principally Disrupts NREM Sleep Brain Activity

Given the absence of gross phenotypes in sleep and circadian behaviour, we next assessed brain activity by applying a Fast Fourier Transform (FFT) to the EEG data. However, *Syngap1^+/-^* mice are known to present interictal spikes (IIS)^48,49^ that could interfere with this analysis, thus, we applied a procedure to identify them (**Figure S2a-c**). Indeed, IIS were increased in *Syngap1*^+/-^ mice (**Figure S2d**), particularly during NREM sleep (**Figure S2e,f)**. Therefore, detected IIS were removed prior to applying FFT to the data.

Analysis of the power spectrums obtained by FFT identified a slowing of the theta peak during wakefulness in *Syngap1*^+/-^ mice (**Figure S3a**). However, this was specific of the electrodes placed above hippocampi (hippocampal derivations), since no differences were observed in the parietal electrode. Regarding sleep, no significant differences were found in the power spectrums of REM sleep between genotypes (**Figure S3b**). Contrarily, we observed robust EEG power alterations in NREM sleep, as these affected all derivations (**Figure 2a**). More precisely, NREM sleep alterations in *Syngap1*^+/-^ animals included an increase in delta (δ, 1-4 Hz) and a decrease in sigma (σ, 9–16 Hz) power, two hallmark rhythms of NREM sleep.

**Figure 2.**
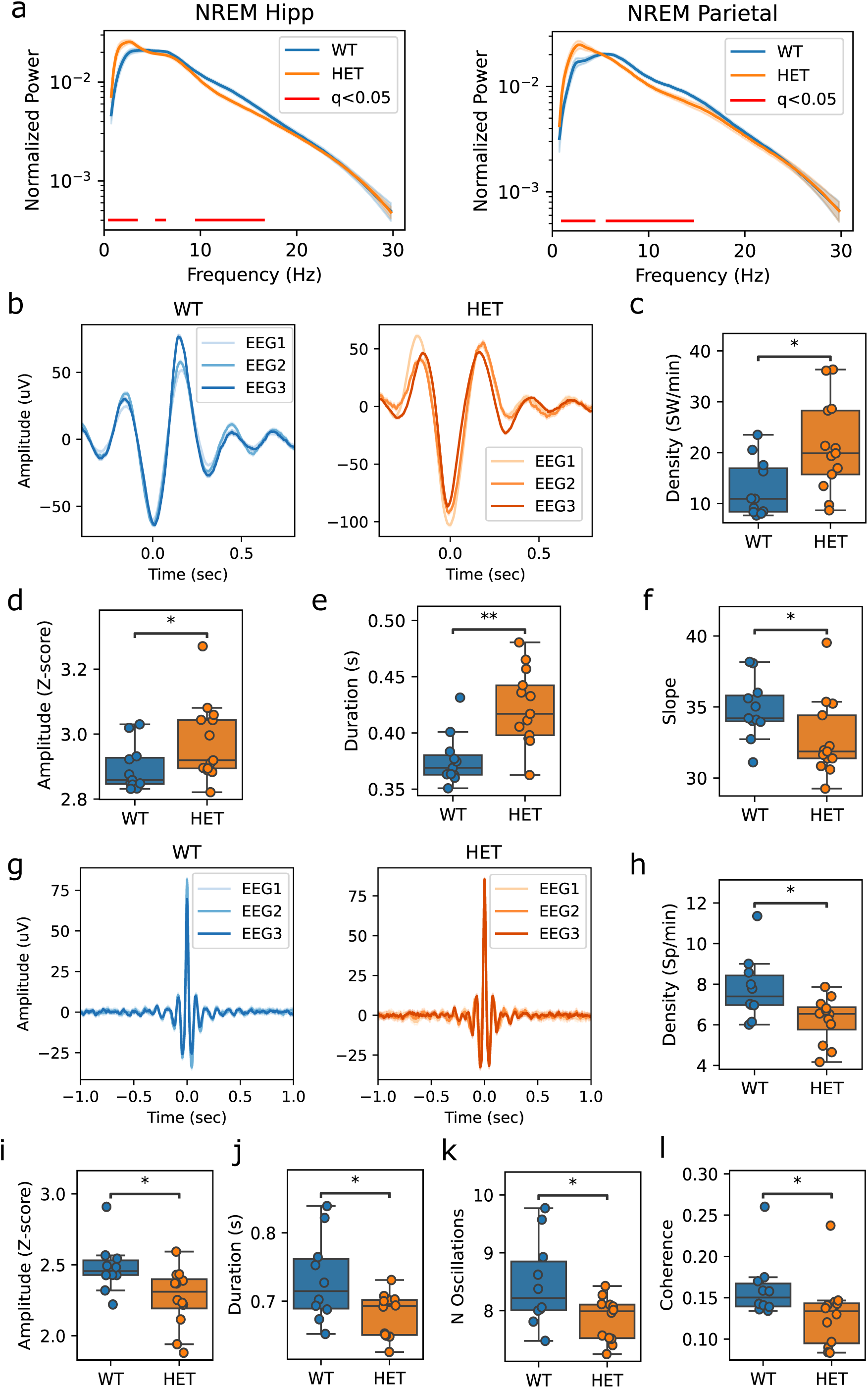
Altered delta and sigma power reflect alterations in slow-waves and spindles in *Syngap*1^+/-^ mice. **(a)** Normalized power spectra during NREM sleep after excluding IIS epochs for both WT (n=11, blue) and HET (n=13, orange) animals in hippocampal channels (left) and the parietal channel (right). Red lines below the traces indicate frequencies with significant differences (multiple independent t-test with Benjamini-Hochberg (BH) FDR correction, q < 0.05). The shaded areas around the line plots represent the SEM. **(b)** Average waveform of detected slow-waves across the three EEG channels (EEG1–3) in representative WT (left, blue) and HET (right, orange) animals. Traces represent the mean ± SEM. Traces are aligned to the main negative peak (time 0). **(c)** Slow wave density (SW/min) during NREM sleep for WT (n=11) and HET (n=13) animals. Dots represent the total number of temporally non-overlapping events across all channels for each animal, normalized to the total artifact-free NREM time. Statistics: Mann-Whitney, p=0.018. **(d)** Average Z-score normalized peak-to-peak amplitude of detected slow-waves per animal. Statistics: Mann-Whitney, p=0.049. **(e)** Mean duration of slow-waves in WT (blue) and HET (orange) animals. Statistics: Mann-Whitney, p=0.002. **(f)** Mean Z-score normalized slope (SD/ms) of slow-waves. Slope was calculated as the peak-to-trough amplitude divided by time between the negative peak and the next 0 crossing. Statistics: Mann-Whitney, p=0.049. **(g)** Average waveform of detected spindles in a representative WT mouse (left) and HET mouse (right) across three EEG channels. Each trace represents the average signal aligned to the spindle peak (time = 0 s). **(h)** Spindle density (sp/minute) during NREM sleep for WT (n=10) and HET (n=11) mice. Dots represent the total number of temporally non-overlapping events across all channels for each animal, normalized to the total artifact-free NREM time. Statistics: independent t-test, p=0.015. **(i)** Mean Z-score normalized amplitude of spindles per animal. Statistics: independent t-test, p=0.02. **(j)** Mean spindle duration (in seconds) calculated across spindles per animal. Statistics: independent t-test, p=0.021. **(k)** Mean number of oscillatory cycles per spindle calculated across spindles per animal. Statistics: independent t-test, p=0.023. **(l)** Mean spindle coherence across EEG channels. Coherence was measured as the mean recall between each pair of EEG channels. Statistics: Mann-Whitney, p=0.018. Boxplots show median, interquartile range (IQR), and 1.5×IQR whiskers; dots represent individual animals.

### Increased Slow-Waves and Reduced Spindles Underpin NREM Sleep Brain Activity Alterations

During NREM sleep, delta and sigma powers reflect the occurrence of slow-waves (SW) and spindles, respectively. In addition, delta power follows a temporal pattern that closely mirrors homeostatic sleep pressure^50–53^, whereas sigma power is regulated in a circadian manner, presenting a peak in the light-dark transition (ZT12)^54,55^. Therefore, the observed power alterations in *Syngap1*^+/-^ mice could arise from differences in SW/spindles, from an impairment of the δ /σ temporal regulation or both.

The analysis of SWs (**Figure 2b**) revealed an increased density of these events in *Syngap1*^+/-^ mice (**Figure 2c**). Furthermore, SWs also presented morphological differences, having higher amplitude and duration (**Figure 2d,e**), and reduced slope (**Figure 2f**). To study the homeostatic regulation of delta power, we fitted a mathematical model of sleep pressure to the temporal pattern of delta power (see Methods). The fitted trajectories closely tracked empirical δ in both genotypes (**Figure S4a-c**). Importantly, the estimated time constants for build-up (Ti; **Figure S4d**) and dissipation (Td; **Figure S4e**) of sleep pressure were similar between genotypes. Together, these data indicate that the increased delta power found in *Syngap1*^+/-^ mice stems from a higher amount and amplitude of SWs, not from differences in the homeostatic regulation of the δ power.

Spindle detection (**Figure 2g**) revealed a reduced density of these events in heterozygous mice (**Figure 2h**). This was accompanied by a decrease in their amplitude, duration and the number of cycles within individual spindles (**Figure 2i-k**). In addition, we also found decreased coherence of spindles between EEG channels, meaning that they are less coordinated across electrodes (**Figure 2l**). Regarding the temporal regulation of σ, both genotypes showed a clear time modulation with similar amplitude and a peak occurring near ZT12 (**Figure S4f-i**). Together, these results show that the sigma power decrease in *Syngap1*^+/-^ mice is driven by a reduction of spindle density, while its circadian modulation is preserved.

Finally, we have previously shown that in *Syngap1*^+/-^ mice IIS mostly appear during NREM sleep. Thus, we asked whether alterations in SWs or spindles were related to this abnormal activity. Interestingly, while no correlation was found between SWs and IIS (**Figure S3c**), a significant negative correlation was found for spindles (**Figure S3d**), suggesting that the epileptic activity in *Syngap1*^+/-^ mice is linked to the reduction of spindle density.

### *Syngap1*^+/-^ Mice Present Attenuated Infra-Slow Oscillations and Fewer Microarousals

Spindles are organized into clusters that appear approximately every 50s (∼0.02 Hz). Spindle clustering drive a rhythm of the sigma power that is known as the Infra-Slow Oscillation (ISO)^56–58^. Importantly, this rhythm is imposed by fluctuations in noradrenaline (NA) levels^57–59^, with the ISO peak coinciding with NA troughs. Since we found reduced spindle density in heterozygous mice, we investigated whether their ISO was also disrupted. We first confirmed that the sigma power of WT animals presented a periodic modulation that coincides with spindle clustering (**Figure 3a**). In contrast, these clusters were not well organised in *Syngap1*^+/-^ mice, resulting in a poor definition of the ISO in these animals. Indeed, FFT of sigma power temporal profile revealed a robust peak between 0.01–0.04 Hz for WTs that was significantly attenuated in our model of SYNGAP1-RD (**Figure 3b**).

**Figure 3.**
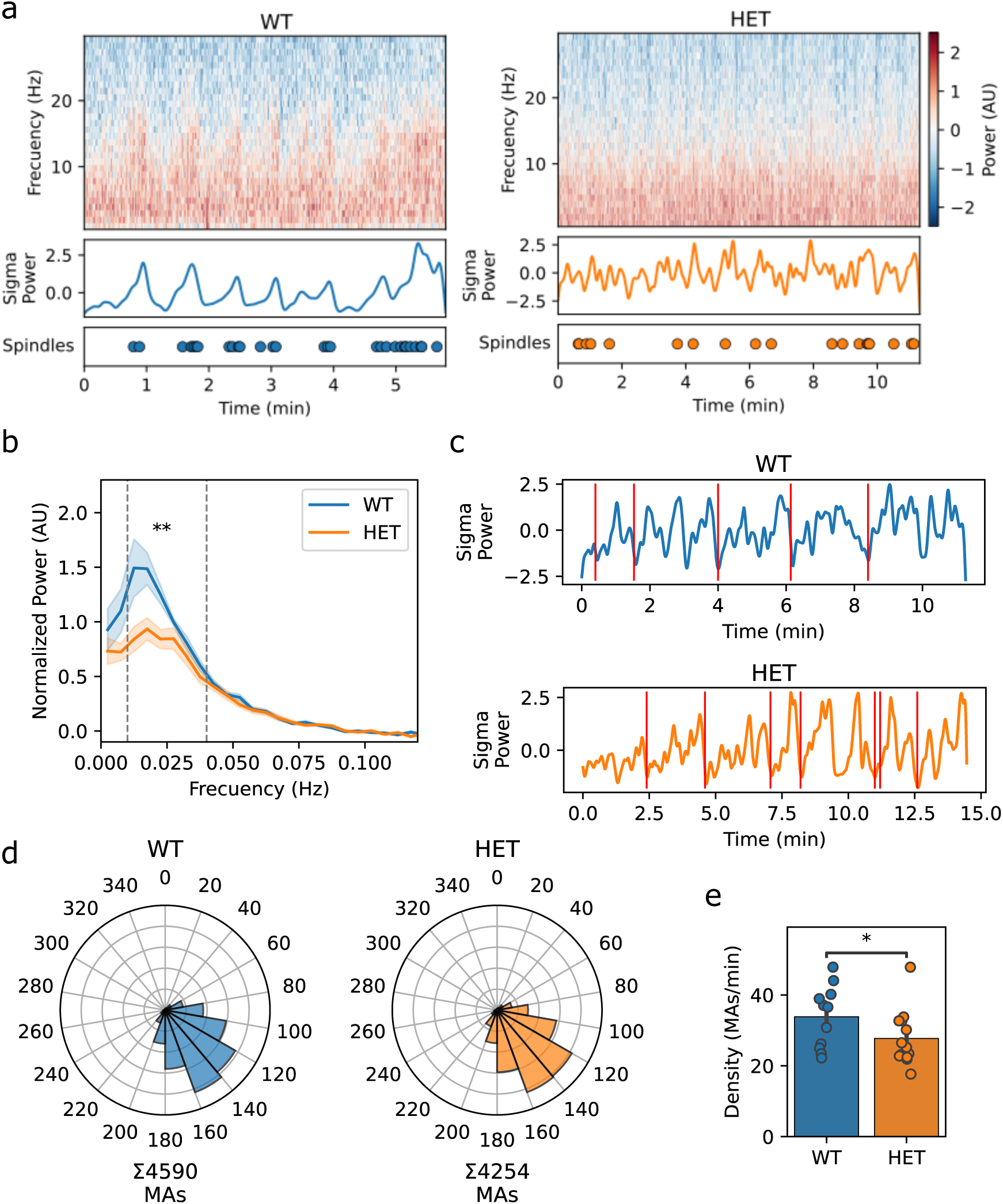
Attenuation of the Infra-Slow Oscillation of Sigma Power in *Syngap*1^+/-^ mice. **(a)** Representative example of a NREM episode from WT mouse (left) and a HET mouse (right). The upper panels show time–frequency spectrograms (Short-Time Fourier Transform; 1–30 Hz), with power z-scored across time and frequency. The middle panels display the slow fluctuations of sigma power (9–16 Hz) obtained with a Morlet wavelet and low-pass filtered at 0.025 Hz to isolate infra-slow dynamics. The bottom panels show spindles detected during the represented episode. **(b)** Group-averaged normalized power spectra of the sigma signal for WT and HET mice after baseline subtraction. Baseline power was estimated from the 0.08–0.12 Hz plateau and subtracted from all frequency bins^58^. The spectra were computed using the FFT of sigma power traces within NREM episodes longer than 96 seconds. Shaded areas represent SEM. Dashed vertical lines indicate the 0.01–0.04 Hz frequency range of interest. For statistical comparison the mean power within the 0.01–0.04 Hz band was calculated for each animal and compared between genotypes with an independent t-test, p = 0.005. **(c)** Examples of a NREM episodes from a WT (top) and a HET mouse (bottom). The slow sigma power trace is overlaid with vertical red bars marking microarousal (MA) onset. **(d)** Circular histograms (20° bins) of the MA phase relative to the σ infra-slow cycle for WT (left, n = 4590 events in 11 animals) and HET (right, n = 4254 events in 13 animals). Zero degrees indicates the σ peak and 180° the trough. Both distributions deviate strongly from uniformity (Rayleigh test: WT p < 10⁻^15^, HET p < 10⁻^15^) and cluster around the descending phase / trough of the oscillation. **(e)** Mean density of MAs (events per minute of NREM) per mouse. Bars represent mean ± SEM; dots are individual animals. Statistical comparison: independent t-test, p=0.016. An outlier was detected through the Z-score method (>2.5 SD) and excluded from the analysis, WT n=11, HET n=12.

Beyond spindle clustering, NA fluctuations critically shape NREM sleep microarchitecture, being linked to the onset of microarousals (MA), which occur in NA peaks and, thus, in ISO troughs^56,60,61^. Therefore, we next quantified the phase relationship between the temporal profile of sigma power and the appearance of MAs (**Figure 3c–d**). MA timing was tightly phase-locked to the ISO in both genotypes near the σ trough. Despite preserved phase-locking, *Syngap1*^+/-^ mice showed a significant reduction in MA density (**Figure 3e**), which is consistent with their attenuated ISO. Overall, the ISO impairment combined with the MA reduction suggest a dysfunction in noradrenergic signalling during NREM sleep in *Syngap1*^+/-^ mice.

Taken together, these findings show that not only spindle density is reduced but their temporal organization is also disrupted. Combined with the alterations in SWs, our results point to an important reorganization of NREM brain activity in *Syngap1*^+/-^ mice, with no major changes in sleep-wake organization and circadian behaviour.

### Disorganization of Daily Synaptic Proteome Dynamics in *Syngap1*^+/-^ Mice

Since the synaptic proteome undergoes a daily remodelling during sleep/wake cycles^18^ and *Syngap1*^+/-^ mice present alterations in NREM sleep, we aimed to investigate if proteome dynamics were affected in these mice. To achieve this, we conducted a proteomics analysis of cortical synaptic fractions collected every 4h along the day (ZT0, ZT4, ZT8, ZT12, ZT16 and ZT20; **Figure 4a**), from WT and *Syngap1*^+/-^ mice, identifying a total of 5,112 proteins (**Table S1**). As previously reported^62^, these synaptic fractions contain proteins spanning all major sub-synaptic compartments (**Figure 4b**), offering a comprehensive view of the synaptic proteome. In parallel, homogenates used to generate these synaptic fractions were processed using the same proteomics workflow (**Table S1**). By comparing these datasets, we aimed to determine whether temporal changes in synaptic protein abundance are synapse-specific or if, instead, they reflect broader, tissue-wide dynamics.

**Figure 4.**
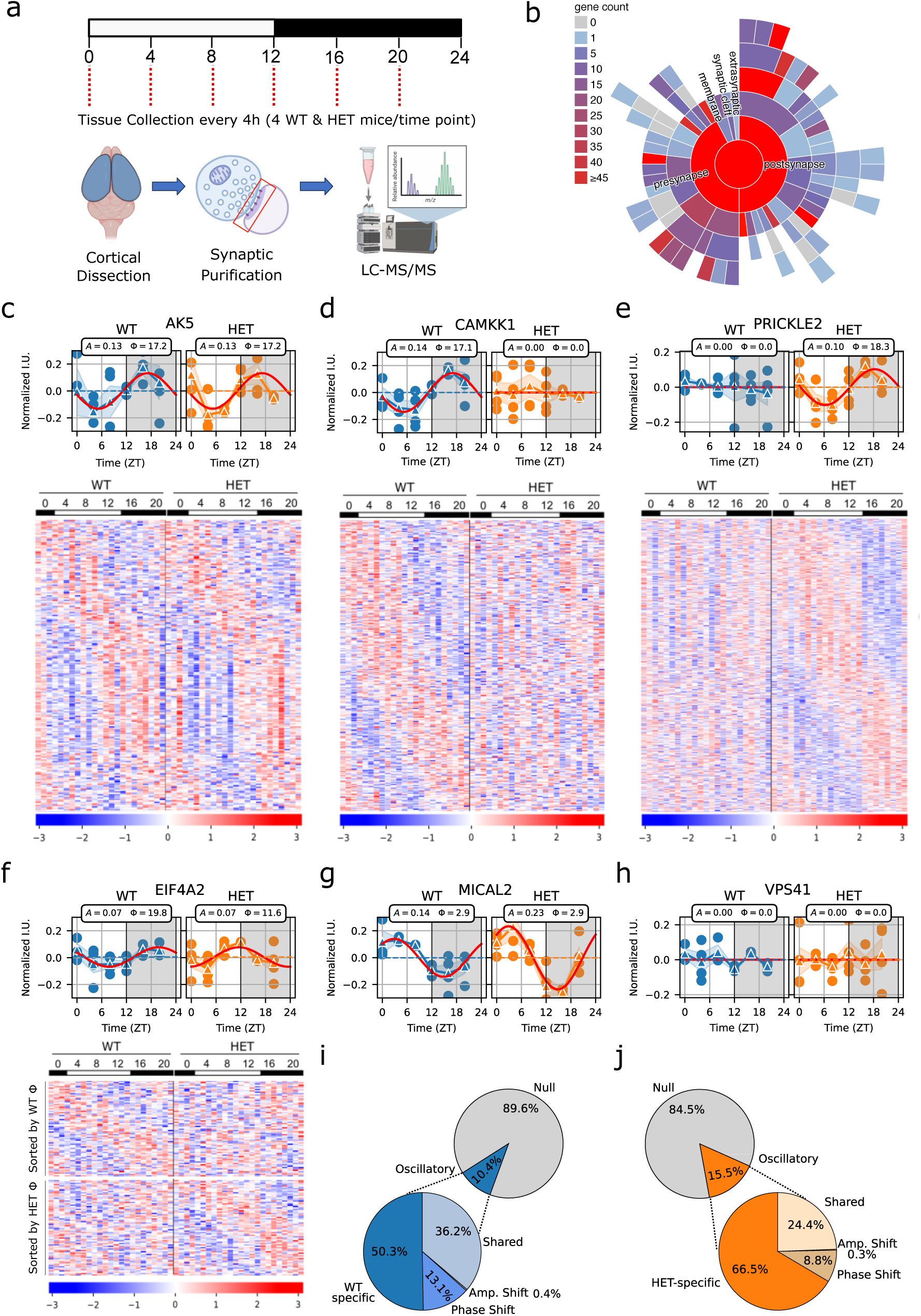
Extensive dysregulation of Homogenate and Synaptic proteome dynamics in *Syngap*1^+/-^ mice. **(a)** Schematic representation of the mass spectrometry-based proteomics workflow used in this study. Cortical samples from WT and *Syngap1*^+/-^ (HET) mice were collected at 6 different Zeitgeber Times (ZT). Synaptic fractions were isolated from each individual animal. These fractions were finally profiled by mass spectrometry. **(b)** Distribution of synaptic proteins across sub-synaptic compartments as defined by SynGO CCs^122^. **(c)** Top, expression abundance along the day of an example protein (AK5) classified in the shared-regulation group. Dots represent individual data points, while triangles connected with a solid blue (WT) or orange (HET) line represent the mean values for each time point. The shaded coloured region surrounding the mean lines represents the SEM. The horizontal dashed represents the baseline (C) parameter. The solid red line represents the fitted cosine function. Shaded grey areas indicate the dark period of the light/dark cycle. The amplitude (A) and phase (Φ) of each protein are shown in the inset. Bottom, heatmap displaying normalized expression of all proteins from the shared-regulation group in both genotypes. Z-scored protein abundances are color-coded, with red indicating higher expression and blue indicating lower expression. Proteins (rows) were sorted by their phase. **(d)** Top, expression abundance along the day of an example protein (CAMKK1) classified in the WT-specific model. Bottom, heatmap displaying normalized expression of all proteins from the WT-specific group in both genotypes. **(e)** Top, expression abundance along the day of an example protein (PRICKLE2) classified in the HET-specific group. Bottom, heatmap displaying normalized expression of all proteins from the HET-specific group in both genotypes. **(f)** Top, expression abundance along the day of an example protein (EIF4A2) classified in the phase-shifted group. Bottom, heatmap displaying normalized expression of all proteins from the phase-specific group in both genotypes. **(g)** Expression abundance along the day of an example protein (MICAL2) classified in the amplitude-shifted group. **(h)** Expression abundance along the day of an example protein (VPS41) falling in the null model. **(i)** Pie charts presenting the proportion of proteins that best fit each model in WT synapses. No proteins were found in the model of amplitude and phase shifting. **(j)** Pie charts presenting the proportion of proteins that best fit each model in and HET synapses. No proteins were found in the model of amplitude and phase shifting.

To identify proteins with daily abundance changes (i.e., cycling proteins) we applied a model selection framework^63^ (see methods, **Table S2**). This includes 7 cosine-based models encompassing all possible expression profiles between genotypes: a shared-cycling model (identifying proteins with the same rhythmic expression pattern in both genotypes; Synapses **Figure 4c**; Homogenates **Figure S5a**;), two genotype-specific models (identifying proteins with rhythmic expression pattern only in one genotype, Synapses **Figure 4d,e**; Homogenates **Figure S5b,c**), three models in which a protein cycles in both genotypes but with shifted amplitude and/or phase (phase being the time of maximal abundance; Synapses **Figures 4f,g**; Homogenates **Figure S5d**) and, finally, a null model, which captures proteins that do not cycle (**Figure 4h**).

In WT mice, approximately 10% (533 proteins) of the synaptic proteome exhibited a cycling behaviour, whereas this proportion increased to 15% in *Syngap1*^+/-^ mice (**Figure 4i,j**). More importantly, only 193 synaptic proteins presented a similar regulation in both genotypes (**Table S2**). Indeed, two-thirds of synaptic proteins cycling in WT, either lost or changed their temporal expression in HET mice (**Figure 4i,j**). Moreover, *Syngap1*^+/-^mice had 526 newly cycling proteins in synaptic fractions (**Table S2**). Noticeably, in WT samples, only a small subset of proteins cycled in both homogenates and synapses (**Figure S5e**) and, of those, few showed similar phases (**Figure S5f**). The same scenario was found for HET samples (**Figure S5g,h**). These differences manifest that homogenate and synaptic fractions largely capture different functional pools of the same proteins and that expression patterns found in synaptic fractions are highly specific to this compartment.

To clarify if genotype-specific proteins reflect robust biological differences, we ranked all proteins by their goodness-of-fit (R²) to a cosine model and partitioned them into deciles. Notably, proteins identified as WT-specific cyclers were significantly enriched in the lowest R² deciles of the *Syngap1*^+/-^ rank (**Figure S6a,b**). Similarly, cycling proteins specific to heterozygous mice were significantly enriched in the lowest R² deciles of the WT rank (**Figure S6c,d**). In contrast, proteins assigned to the shared-cycling, phase-shift and amplitude-shift models exhibit a significant enrichment within the highest R² deciles in both genotypes (**Figure S6e-h**). These results indicate that proteins classified by the genotype-specific models reflect robust loss or gain of cycling proteins in *Syngap1*^+/-^mice.

### Synaptic Functions in Wild-Type Mice Are Organized Sequentially Along the Day

We first wanted to investigate how regulation of synaptic protein abundance shapes synaptic function along the day in wild-type mice (**Figure 5a**). To do this, we performed an enrichment analysis of signalling pathways and Gene Ontology (GO) terms using all proteins with a temporal regulation. However, we argued that for a given term to be considered as time-regulated most of its constituent proteins should cycle in a synchronized manner. Thus, we introduced the notion of temporal coherence in this analysis (see methods). In addition, temporally coherent terms (**Table S3**) were grouped into the following 9 broader categories, termed Synaptic Functional Modules (SFM): Signal Transduction, Translation, Metabolism, Protein Stability, Intracellular Trafficking, Endocytosis, Synaptic Vesicle, Synaptic Assembly and Synaptic Stabilization.

**Figure 5.**
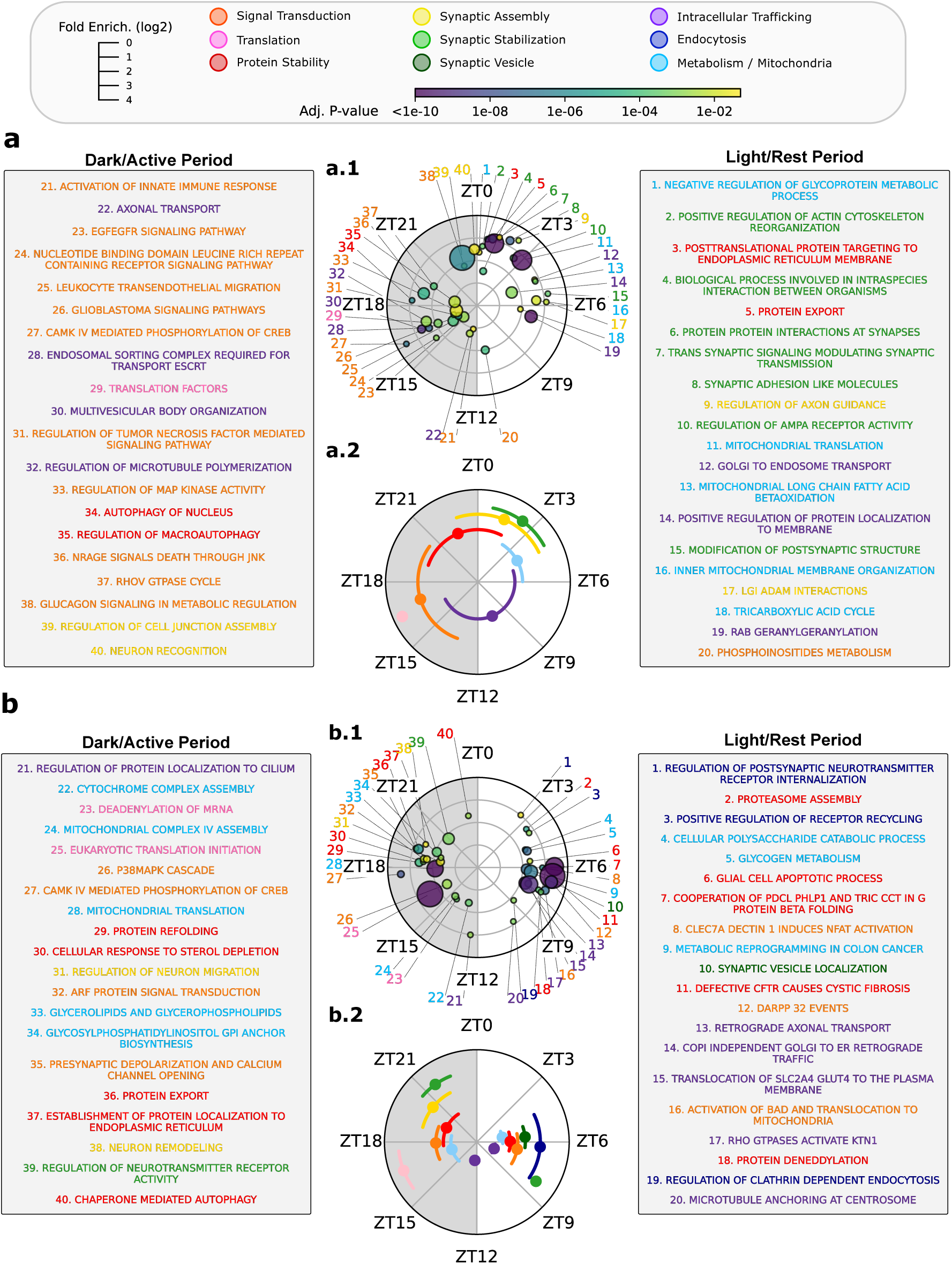
Temporal coherence analysis of enriched pathways reveals altered distribution of synaptic functions along the day. **(a)** Synaptic function organization along the day in WT animals. (a1) Scatter plot displaying the top 20 time-coherent enriched pathways and GOBP for the dark/active period (left semicircle) and the light/rest period (right semicircle). Each dot represents a term, with the angular position representing the mean phase of its constituent proteins. The distance from the centre (radial depth) reflects fold enrichment (log2-transformed), dot size reflects the number of proteins in the term and the colour of dots indicates the adjusted p-value according to the presented scale. Terms were classified in different Synaptic Functional Modules (SFM) and their numeric label and term description (right and left boxes) were colour-coded as stated in the upper legend. (a2) Summary representation of SFMs along the day. Each colour represents a SFM (see legend in the top) and its position in the graph represents the mean (dot) +/- circular std (line) calculated using the phase of all terms included in such category. **(b)** Synaptic function organization along the day in HET animals. (b1) Scatter plots displaying the top 20 time-coherent enriched pathways and GOBP for the dark/active period (left semicircle) and the light/rest period (right semicircle). (b2) Summary representation of SFMs along the day. Light and dark periods were treated separately to better show the biphasic distribution of their temporal organization.

The dark/active period (ZT12–ZT24) was characterised by three major SFMs: Signal Transduction, Translation and Protein Stability (**Figure 5a2**). Signal Transduction pathways, with terms related to regulation of MAPK, calmodulin-dependent kinase activity, or PKC biology (**Table S3**). Notably, of the 202 kinases present in our synaptic proteome 30 cycled in a daily fashion, with 26 of them peaking in the dark (**Table S2**). Examples include Camk2a, Camk2g, Pkcg or Sik3. Noteworthy, Sik3, which has been consistently implicated in sleep regulation^64–68^, cycled with a maximum at ZT19 in WT synapses.

Protein Translation control was also activated early in the dark phase, with multiple initiation factors (EIF1AX, EIF3E, EIF3G, EIF4A1, EIF4A2) peaking at ∼ZT16. Subsequently, in the second half of this period, several processes related to Protein Stability emerged. Proteins relevant to autophagy (ATG5, ULK2, WDR45) peaked at ∼ZT18, while terms related to the unfolded protein response (ER/UPR), known to be upregulated by extended wakefulness^69–71^, emerged near the dark-to-light transition (**Figure 5a1, Table S3**).

Interestingly, terms from the Synaptic Assembly module appeared at the end of the dark phase and spanned into the early-light period. These include “Regulation of cell junction assembly” at ZT23, the term with the largest number of proteins (23), as well as others like “Neuron recognition” or “Regulation of axon guidance”. These terms contain key proteins for synaptic formation, such as neuroligins (NLGN2, NLGN3), Bai1 (ADGRB1), NRP1, NEGR1 or FLRT3; as well as neurotrophic receptors EPHA7, SLITRK2, SLITRK4, TrkC (NTRK3) and their ligands, SLIT1 and EFNB3 (**Table S3**). Remarkably, during the early-light period terms related to the Synaptic Stabilization module were prominent, including functions important for postsynaptic organization or for the localization and activity of neurotransmitter receptors. These terms included core PSD scaffolds (DLG3, DLGAP3, HOMER2, SHANK2), glutamate receptors (GRIA4, GRIN2B, GRM5), their auxiliary subunits (CACNG2/7; NETO1) and actin regulators (BAIAP2, RHOA).

The first half of the light period was also characterized by the emergence of pathways related to metabolism and mitochondrial function (e.g., β-oxidation or TCA cycle). Strikingly, proteins involved in mitochondrial translation, including many ribosome components and two translation initiation factors, showed maximal expression at ZT3, suggesting intense mitochondrial protein synthesis at this time. Finally, starting at the second half of the light period and spanning into the dark one, synaptic function was dominated by the Intracellular Trafficking module (e.g. Golgi vesicle biogenesis, microtubule motor activity, axonal transport).

Overall, in WT animals, SFMs were clearly segregated along the day, with little overlap between them. Remarkably, our results suggest that synaptic functions are dynamically adjusted to support information processing during the active period, resulting in synaptic remodeling or formation towards the dark-to-light transition. These events would culminate, during the first hours of the rest phase, in a process of synaptic stabilization (**Figure 5a2**).

### Marked Alterations in Synaptic Functions During the Light/Rest Period in *Syngap1*^+/-^ Mice

In *Syngap1*^+/-^ mice synaptic function organization in the dark/active period was overall well preserved (**Figure 5b** and **Table S3**). As in WT mice, this period was mostly characterized by terms related to Signal Transduction (e.g., CaMKIV–CREB, p38 MAPK, kinase activator activity), Protein Translation and Protein Stability (e.g., folding/stress-response terms; chaperone-mediated autophagy). For instance, 16 of the 26 kinases peaking in the dark in WT animals also did so in HET synapses (e.g. CAMK2A, PRKCG or ROCK2), although proteins with gained regulations also contributed to Signal Transduction functions (e.g. CAMK2B, PLCD3, DAP2IP or TIAM1). Regarding Protein Translation, it is noteworthy that *Syngap1*^+/-^ synapses not only presented increased levels of translation initiation factors, as observed in WTs, but also many ribosomal subunits peaked at ZT17. Finally, as in WTs, terms from the Synaptic Assembly module (e.g., neuron remodelling, focal adhesion assembly, positive regulation of cell-junction assembly) rose towards the end of the dark cycle. These contained shared cycling proteins such as NRP1, NLGN3, FLRT3 but also gained ones as EFNB2, PTPRF or RELN.

Nevertheless, some differences were also observed during the dark/active period. For instance, terms from the Synaptic Stabilization module, which in WT mice appear in the early light/rest period, were found in the dark. These included terms such as ‘ionotropic glutamate receptor activity’, ‘neurotransmitter receptor complex’, or ‘regulation of neurotransmitter receptor activity’, containing proteins like GRIK2, GRIN2D, GRID1, DLGAP4 or SHISA6. Similarly, a cluster of metabolism and mitochondrial-related terms, which in WTs are characteristic of the light period, also appeared during darkness. These involved cellular respiration and mitochondrial protein synthesis, meaning that the prominent peak of mitochondrial ribosome components appearing around ZT3 in WT samples was shifted close to 12 hours in heterozygous mice.

In contrast to the better conservation of SFMs found in the dark/active period, more pronounced divergences emerged in the light/rest one. Most noticeably, terms related to Synaptic Stabilization in the early light phase were absent from *Syngap1*^+/-^ synapses. Instead, we observed multiple terms from two related SFMs unique to HET samples: Endocytosis and Synaptic Vesicles. These SFMs were related to the internalization of neurotransmitter receptors and to clathrin-mediated endocytosis, which were driven by the cycling of clathrin subunits (CLTA and CLTC), clathrin adaptors (PICALM and SNAP91), cargo selection proteins (e.g. ARRB2 or DAB2) and many synaptic vesicle constituents (e.g., SYN1, SYP, SVOP, SCNA). In addition, while in WTs Signal Transduction were mostly confined to the dark period, in HET synapses this module was also abundant during the light. However, these Signal Transduction terms were clearly different from those in the dark, including phosphatases and kinases related to synaptic depression (PPP3CA, PPP3CB, GSK3B and CDK5)^72–75^. Together with the appearance of endocytic processes, these results suggest a process of synaptic weakening in *Syngap1*^+/-^ synapses during the light/rest period.

Further differences in synaptic functions from the light/rest period include a strong presence of Protein Stability terms, which in WTs appeared mainly confined to the dark. Indeed, a very large number of components from the proteasome (19/33) and the Chaperonin Containing TCP-1 (CCT) complex (8/9) cycled coherently in *Syngap1*^+/-^synapses with maximal abundances around ZT6. On the other hand, two SFMs characteristic of the light were preserved in HET mice: Metabolism/Mitochondria (e.g., TCA, glycolysis/gluconeogenesis) and Intracellular Trafficking (e.g., axonal transport, dynein heavy-chain binding).

Overall, the temporal organization of SFMs in *Syngap1*^+/-^ mice was importantly altered, showing increased temporal overlap and clustering either in the mid-light or mid-dark period (**Figure 5b2**). However, we found the identity of SFMs to be better preserved during the dark period than in the light one. Our results suggest that, as in WTs, synapses are primed for information processing in the active phase, leading to synaptic formation. However, the synaptic stabilization occurring during early-rest would be switched for a process of synaptic downscaling.

### NREM Sleep Is the Main Driver of Synaptic Proteome Dysregulation in *Syngap1*^+/-^ Mice

Since the disorganization of synaptic functions in HET mice was most pronounced during the light period, we examined whether genotype-specific cycling proteins preferentially peaked during this circadian window. We therefore analysed the distribution of phases among shared and genotype-specific cycling proteins. Interestingly, phases within each group were not randomly distributed across the day but, instead, clustered at specific times (**Figure 6a-c**). Shared cycling proteins tend to peak at ZT19 (**Figure 6a**), proteins with lost regulation in HET mice (i.e WT-specific) do so around ZT4 (**Figure 6b**) and HET-gained cycling proteins exhibit a biphasic distribution, with peaks at the middle of the light (ZT6.5) and dark (ZT18.5) periods (**Figure 6c**). Thus, shared cycling proteins tend to peak at the dark/active period (**Figure 6a**), where synaptic functions are better preserved (**Figure 5b**), while proteins from the genotype-specific models tend to do so in the light/rest period, which presented marked functional differences (**Figure 5b**).

**Figure 6.**
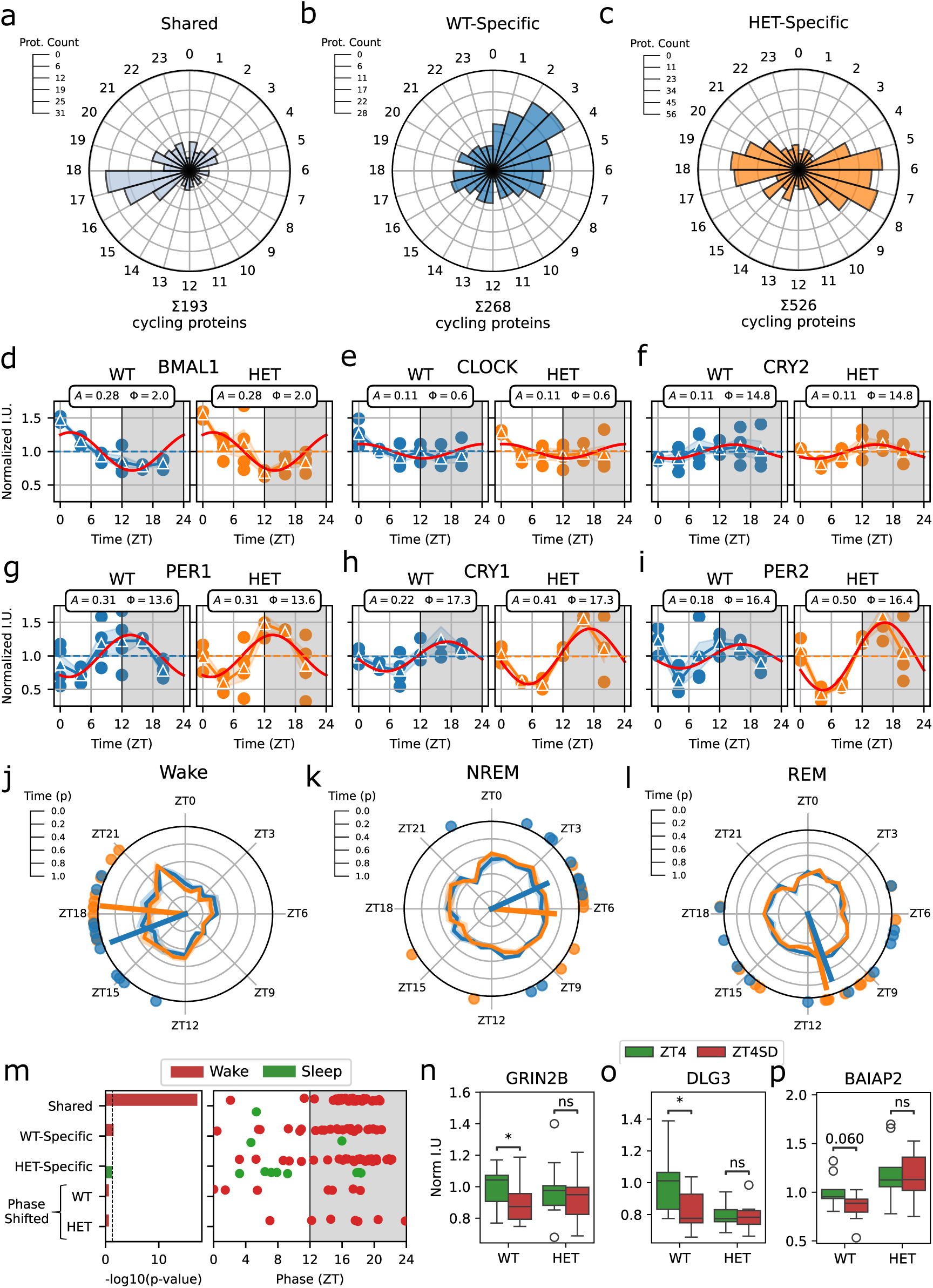
Synaptic proteome dynamics are driven by sleep-wake cycles. **(a-c)** Circular histogram showing the phase distribution of common rhythmic proteins (a), WT-specific oscillatory proteins (b) or HET-specific oscillatory proteins (c). **(d-i)** Expression abundance along the day of circadian clock genes: (d) BMAL1, (e) CLOCK, (f) CRY2, (g) PER1, (h) CRY1 and (i) PER2. Dots represent individual data points, while triangles connected with a solid blue (WT) or orange (HET) line represent the mean values for each time point. The shaded region surrounding the mean lines represents the SEM. The horizontal dashed represents the baseline (C) parameter. The solid red line represents the fitted cosine function. Shaded grey areas indicate the dark period of the light/dark cycle. The amplitude (A) and phase (Φ) of each protein are shown in the inset. **(j-l)** Circular distribution of Wake (j), NREM (k) and REM sleep (l) states. Blue (WT) and orange (HET) lines represent the mean proportion of time in the state per hour of recording. Shaded regions surrounding the mean represent SEM. Dots represent the circular mean (μ) of the distribution of individual animals. Arrows represent the mean resultant vector for each genotype, where the length of the vector (r) represents the data aggregation. Wake: WT μ=16.6 and r=0.89, Rayleigh test p=1.69e-5; HET μ=18.4 and r=0.95, Rayleigh test p=1.05e-7. NREM: WT μ=4.4 and r=0.69, Rayleigh test p=0.003; HET μ=6.3 and r=0.72, Rayleigh test p=0.005. REM: WT μ=10.7 and r=0.44, Rayleigh test p=0.12; HET μ=11.1 and r=0.64, Rayleigh test p=0.003. **(m)** Comparison of cycling proteins in our study with those previously published sleep/wake regulated proteins. Left panel: Enrichment of the proteins in each of our models in proteins found upregulated by Wake (red) or Sleep (green) in a previous publication^16^. Right panel: Phase distribution of overlapping proteins between each of our models and Wake- or Sleep-upregulated proteins. Each dot represents an individual protein. **(n-p)** Immunoblot quantification of (n) GRIN2B, (o) DLG3 and (p) BAIAP2 in WT and HET undisturbed (Green, ZT4) or sleep-deprived (Red, ZT4SD) animals collected at ZT4. Boxplots show median, interquartile range (IQR), and 1.5×IQR whiskers; empty dots represent individual animals outside the IQR. Statistics: independent t-test. GRIN2B: WT p= 0.037, ZT4 n=12, ZT4SD n=12; HET p=0.45, ZT4 n=12, ZT4SD n=12. DLG3: WT p= 0.037, ZT4 n=9, ZT4SD n=9; HET p=0.86, ZT4 n=9, ZT4SD n=9. BAIAP2: WT p= 0.06, ZT4 n=9, ZT4SD n=9; HET p=0.75, ZT4 n=9, ZT4SD n=9.

The non-uniform distribution of protein phases within each group indicates that they might be regulated by a common driver. Two plausible candidates would be circadian rhythms and sleep/wake cycles^18^. To explore if a circadian dysregulation existed at the molecular level in heterozygous animals, despite their preserved circadian behaviour (**Figure 1, S1 and S3**), we analysed the daily gene expression pattern of six circadian clock components in the same cortical samples used for proteomics. In line with our behavioural data, we found cycling patterns of these genes to be well preserved in HET mice (**Figure 6d-i**), further arguing against circadian rhythms acting as the main driver of synaptic proteome dysregulation.

We thus explored the contribution of sleep/wake cycles to synaptic proteome dynamics by looking at the temporal distribution of wakefulness, NREM and REM in our EEG data. Wakefulness distribution is centred around ZT16 in WTs and ZT18 in HET mice (**Figure 6j**), close to the mean phase of shared-cycling proteins (**Figure 6a**), suggesting they could be upregulated by wakefulness. Regarding sleep, REM shows a distribution centred around ZT11 in both genotypes (**Figure 6l**), yet this does not coincide with the mean phase of any protein cycling group. Indicating that REM sleep is not a major driver for any of these protein groups. On the other hand, NREM sleep in WTs is centred around ZT4 (**Figure 6k**), matching the mean phase of WT-specific cycling proteins (**Figure 6b**). In HETs, this centre is at ZT6 (**Figure 6k**), also matching the main peak of HET-specific cycling proteins (**Figure 6c**). This would support that genotype-specific cycling proteins are influenced by NREM sleep.

To further investigate the connection between synaptic proteome dynamics and sleep/wake cycles, we took advantage of a recently published synaptic proteomics study, which uses a sleep deprivation (SD) paradigm, allowing to unambiguously determine proteins that are regulated by sleep/wake cycles^16^. Importantly, we found a highly significant overlap between our genotype-shared cycling proteins, mostly peaking during the active period, and those upregulated by wakefulness in that study (**Figure 6m**). These results strongly suggest that most proteins with preserved rhythms in HET mice are upregulated by wakefulness. On the contrary, there was little overlap between the sleep regulated proteins and the cycling proteins from any group in our study. Thus, to explore the association between genotype-specific cycling proteins and NREM sleep, we collected cortical tissue from mice at ZT4 either after 4h of SD or normal rest. We immunoblotted synaptic fractions from these mice for GRIN2B, DLG3 and BAIAP2 (**Figure 6n-p**), three WT-specific cycling proteins related to Synaptic Stabilization.

GRIN2B and DLG3 were significantly reduced after SD in WT and BAIAP2 showed a similar non-significant trend (p=0.078). These results indicate that these proteins are upregulated by sleep in WT animals. Of note, SD did not affect the level of these proteins in *Syngap1*^+/-^ mice (**Figure 6n-p**), consistent with their loss of cycling behaviour.

Together, our findings suggest that the greater disruption of synaptic functions occurring during the light period in *Syngap1*^+/-^ mice is linked to the preferential peak of genotype-specific cycling proteins during this period. Importantly, while shared cycling proteins are related to wakefulness, genotype-specific ones would be upregulated by NREM sleep. These findings are in line with NREM sleep being the most affected behavioural state in *Syngap1*^+/-^ mice as observed in the EEG analysis.

## DISCUSSION

While the importance of NREM oscillations on sleep-dependent memory consolidation is well-stablished^3,4,21^, their impact onto the proteomic composition of synapses is ignored. Here, through the integrative analysis of EEG recordings and time-resolved synaptic proteomics in a NDD model, we propose that slow-waves and spindles, two brain oscillations characteristic of NREM sleep, would mediate synaptic proteome remodeling during sleep. Importantly, our results indicate that this sleep-dependent remodeling results in a synaptic stabilization, which could serve as the molecular substrate of memory consolidation occurring during sleep.

In WT animals 10% of the cortical synaptic proteome is subjected to daily expression changes. Consistent with previous findings^18^, these dynamics are strictly synaptic, since changes at the whole cortical level are essentially different. Importantly, this reveals that daily proteome dynamics would be driven by synapse-specific mechanisms. As previously shown^18^, we found these proteomic dynamics to be linked to sleep/wake cycles. Indeed, proteins peaking in the active period tend to do so when wake density is maximal. Furthermore, these proteins increase their levels with sleep deprivation (SD), which is indicative of a wake-driven upregulation. Likewise, proteins peaking in the rest period do so near the time of highest NREM sleep density and decrease upon SD, pointing to a NREM-driven upregulation. Together, these findings suggest that different brain states impose distinct patterns of synaptic stimulation, leading to specific synaptic proteome changes independently of tissue-wide changes.

Remarkably, daily synaptic proteome dynamics result in the temporal segregation of key synaptic functions. Throughout the active period, WT synapses are strongly engaged in signal transduction, in agreement with theoretical frameworks proposing that synaptic signalling increases during wakefulness as a result of experience-dependent neuronal activity^23,76,77^. Simultaneously, the peak of translation initiation factors would support activity-dependent translation. Later in the active period, protein degradation and folding pathways appeared, suggesting that sustained synaptic activity during this period leads to molecular stress. This is in line with prolonged wakefulness activating the unfolded-protein response (UPR)^69–71^.

Towards the dark–light transition, the upregulation of proteins involved in cell-adhesion and synaptic assembly suggests a period of enhanced synaptic remodelling and formation, which extends into the early light phase. This is accompanied by the upregulation of core PSD scaffolds, glutamate receptors and their regulators, which we interpret as a period of synaptic stabilization. The early light/rest period is also characterised by mitochondrial and metabolic events. Especially notable was the coordinated overexpression of mitochondrial ribosome proteins at ZT3, which is indicative of a window for synaptic mitochondria regeneration. Consistently, SD leads to mitochondrial deficits^78–81^, and mitochondrial metabolism contributes to sleep-wake regulation^79,82^. Finally, the second half of the light/rest period presented enrichment of intracellular transport, in line with transcriptomics studies showing that sleep upregulates these processes^83–85^.

Overall, our findings in WT animals support the notion that wakefulness-related signalling prime synapses for later stabilization during sleep^77^. Indeed, we observed that synaptic functions are nicely segregated along the day, supporting information processing during the active period and leading to synaptic remodeling or formation towards the dark-to-light transition. These events would culminate, during the first hours of the rest period, with a process of synaptic stabilization.

Importantly, this orchestrated organization of synaptic functions across the day was severely disrupted in *Syngap1*^+/-^ synapses, with all SFMs showing great temporal overlap. This functional disorganization was more pronounced during the light period, when animals are mostly asleep. Accordingly, WT cycling proteins upregulated by wakefulness were better preserved in *Syngap1*^+/-^ mice than those regulated by sleep, pointing to a sleep-dependent proteomic dysregulation in *Syngap1*^+/-^ mice. However, the amount of wake, NREM and REM sleep was equivalent between genotypes, thus sleep architecture *per se* is unlikely to explain the proteomic differences observed. Sleep alterations in *Syngap1*^+/-^ mice were essentially restricted to the two major oscillations characteristic of NREM sleep, slow-waves, which were enhanced, and spindles, which were reduced and temporally disorganised, suggesting that these oscillations mediate the remodelling of the synaptic proteome.

Indeed, the alterations in NREM oscillations are well suited to explain the proteomic dysregulation found during the rest period, since both have been proposed to play important, yet opposed, roles in synaptic plasticity during sleep^30^. While SWs facilitate synaptic downscaling^24–26^, spindles promote synaptic potentiation trough Ca^2+^ influx into dendrites^27–30^. Therefore, increased SWs combined with reduced density and infra-slow organization of spindles, may underlie a shift from synaptic stabilization towards synaptic weakening. This shift is reflected by the enrichment in *Syngap1*^+/-^ synapses of processes related to endocytosis, neurotransmitter receptor internalization, and kinase/phosphatase signaling associated with LTD and homeostatic downscaling^72–75^. Thus, our findings are in line with the proposed roles for NREM sleep oscillations in synaptic plasticity, suggesting that spindle-rich NREM sleep promotes synaptic potentiation while SW-rich NREM sleep promotes downscaling^30^. In addition, the loss of synaptic stabilization in *Syngap1*^+/-^ mice might affect their capacity to consolidate memories during sleep, impacting their cognitive performance.

Disturbances in sleep are more common in children with NDDs than in their neurotypical peers^12–14^. Furthermore, alterations in NREM oscillations have been reported in monogenic NDDs, including Rett^35,86^, Angelman^36,87,88^, Fragile X^37,89–91^, Phelan–McDermid^92,93^ or Dravet^38,94,95^. Among these alterations, the decrease in spindle density seems to be a common trait, both in human patients and rodent models of this conditions^35,36,92–95,37,38,86–91^. Noticeably, we recently reported a decrease in spindle activity in SYNGAP1-RD patients^33^, what underscores the relevance of our findings in the mouse model. This raises the possibility that dysregulation of synaptic proteome dynamics is a common feature in NDDs coursing with sleep disorders. Further studies directly manipulating specific features of NREM sleep, particularly SWs and spindles, would contribute strengthening the observations linking these oscillations with proteome remodelling. Furthermore, targeting these oscillations may hold therapeutical potential for SYNGAP1-RD and other NDDs, since the recovery of normal synaptic proteome dynamics might result in improved cognitive performance.

Overall, our studies on a NDD-causing gene allows us to propose that NREM oscillations mediate the synaptic proteome remodelling occurring during sleep, which would be linked to the stabilization of synapses and, potentially, to memory consolidation. Furthermore, we reveal a putative pathological mechanism that could be common to neurological conditions involving alterations in NREM oscillations.

## METHODS

### Animal handling

Animal research was done with equal numbers of male and female inbred mice from the conditional *Syngap1* rescue line (Syngap1^+^/^lx-st^, Jackson Laboratories, strain code JAX: #029304) in accordance with national and European legislation (Decret 214/1997 and RD 53/2013). Research procedures were approved by the Ethics Committee on Animal Research from the: i) Institut de Recerca Sant Pau (IR-SANT PAU) ii) Centro Andaluz de Biologia Molecular y Medicina Regenerativa (CABIMER) iii) The Herbert Wertheim UF Scripps Institute for Biomedical Innovation & Technology. These procedures were also approved by the Departament de Territori i Sostenibilitat from the Generalitat de Catalunya (approval reference numbers 9,655 and 164.16). Maintenance and experimental procedures were conducted at the animal facilities of the IR-SANT PAU and CABIMER. Mice were housed at a 12h light/dark cycle, at a temperature of 22 ± 1 °C and a relative humidity of 55% ±10%. Mice had access to fresh water and food ad libitum. We used animals of both sexes and 9-14 weeks of age.

### Mouse Brain Dissection

Mice were culled by cervical dislocation minimizing animal suffering. Mouse head was severed with dissecting scissors and placed onto a glass petri dish with a filter paper soaked with chilled 1x PBS (10 mM Na2HPO4, 1.8 mM KH2PO4, 137 mM NaCl, 2.7 mM KCl, pH 7.4). The skin was removed by a cut through the long axis of the head, using 14.5 cm dissecting scissors, whereas the skull and meninges were removed using 11.5 cm Iris scissors and tissue forceps 1:2 (Thermo Scientific). The brain was then extracted using curved Gerald thumb forceps (VWR International, Darmstadt, Germany), rinsed with chilled PBS, and dissected with two spatulas following Spijker (2011)^96^. Tissue weighted, snap-frozen in liquid nitrogen and stored in a -80°C freezer (Thermo Fisher Scientific).

### Mouse Video Recording and Animal Tracking

Video recordings were conducted to assess circadian behaviour in HET mice. Four cages were recorded simultaneously using an infrared camera (Basler acA640-750um camera, ref. 106748) positioned above the cages, at a sampling rate of 30Hz. Recordings started at Zeitgeber Time 0 (ZT0; 7:00 a.m.) and lasted for 48 hours. Mice were individually placed in clean cages at the start and remained undisturbed throughout. Lighting conditions matched the 12 h light/dark cycle of the animal facility.

Videos were downsampled 60-fold using ffmpeg (https://www.ffmpeg.org/). Animal tracking was performed with DeepLabCut v3.0^97–99^ using the pretrained SuperAnimal-TopViewMouse model^100^. The following body parts were tracked: nose, eyes, ears, head, neck, shoulders, back, hips, and tail. The animal’s centroid was computed from major body parts (excluding the tail), omitting coordinates with a detection likelihood below 0.95. Additionally, to avoid misidentification of body parts, the area covered by the mouse’s body was calculated per time point and whenever it was >100 pixels the centroid position was discarded and filled with the next valid position. Distance travelled was calculated as the pixel difference between consecutive centroid coordinates. Activity during light and dark periods was quantified as total distance travelled per periods and compared using a mixed ANOVA, with Genotype as the between-subject factor and Circadian Phase as the within-subject factor.

### Electrode Implantation and Electroencephalogram (EEG) Recordings

Stereotaxic surgeries and EEG recordings were conducted at CABIMER (Seville, Spain). Mice were anesthetized with isoflurane (2–3% induction, 1–2% maintenance; 0.8 L/min O₂ flow), and eight Teflon-coated silver-nichrome wire electrodes (125 µm diameter; A-M Systems, USA) were bilaterally implanted in the frontal cortex (±1 mm lateral, +1 mm anterior to bregma), parietal cortex (±2 mm lateral, 0 mm anterior), and above the dorsal hippocampus (±2.5 mm lateral, −2.5 mm posterior). Two additional electrodes served as ground (−2.0 mm lateral, −4.8 mm posterior) and reference for both hippocampal electrodes (+2.0 mm lateral, −3.5 mm posterior). Frontal and parietal electrodes were referenced to the contralateral hemisphere. Electrodes were connected to a pin header linked to a signal amplifier (8406-SE4, Pinnacle Technologies) and secured with dental cement, with the entire assembly encased in acrylic resin.

After surgery, mice were allowed to recover for at least 7 days. Recordings were performed on 14-16 weeks-old freely moving animals using a synchronized video-EEG monitoring system (8400-K1-SE4, Pinnacle Technologies) equipped with four channels and managed by Sirenia® Acquisition software. Sampling rate was 250 Hz and preamplification gain was set at 100x. Mice were housed individually in cages and monitored for 24-hour sessions.

### Sleep Architecture Analysis

Raw EEG data were visually inspected using SleepSign (Kissei Comtec, Japan) and classified into Wake, NREM, and REM stages in 4-second epochs. Artifacts were marked and excluded to ensure data quality.

The proportion of time spent in each vigilance state (Wake, NREM, REM) was calculated for the full recording period and normalized by the total duration of the recording. To assess circadian variation, data were binned according to the light/dark phases. For comparisons of sleep patterns over the 24-hour cycle, data were binned by Zeitgeber Time (ZT). Statistical analyses were performed using mixed ANOVA with Genotype as the between-subject factor and Vigilance State, Circadian Period, or ZT as within-subject factors.

Episodes were defined as continuous periods of a given state allowing brief interruptions by other states, with minimum durations of ≥52 s (13 epochs) for Wake/NREM and ≥32 s (8 epochs) for REM. Interruptions ≤4 consecutive epochs (16 s) and <30% of total duration were permitted^27,101^. Episodes were detected using a custom-made Python function. The number of episodes were normalized by total recording time and compared between genotypes using independent t-test. Episode stability was assessed with the lifelines Python package^102^ using Cox regression models accounting for repeated measures and circadian phase and Kaplan–Meier curves for visualization.

### Interictal Spike (IIS) Detection and Quantification

Raw EEG signals were processed using MNE-Python^103^. IIS detection was conducted by adapting previously described methods^104^. Specifically, the find_peaks function from SciPy was applied to both raw and inverted EEG signals to detect positive and negative peaks, respectively. Final detection criteria were as follows: Prominence ≥ 200 µV, Height ≥ 2 × baseline, Width < 200 ms, Inter-spike interval ≥ 100 ms. Baseline was dynamically calculated for each subject and vigilance state as the 99th percentile of the absolute signal amplitude. IIS frequency was quantified as spike density (spikes/hour) for each animal and compared using mixed-design ANOVA with Genotype as the between-subject factor and Stage as the within-subject factor. Post hoc tests were conducted using pairwise comparisons with Holm correction. To visualize the shape of IIS, segments centred on the peak of each spike (± 300 ms) were extracted and averaged to generate mean waveforms per animal, channel, and polarity (positive, negative).

### Spectral Power Analysis

EEG signals were pre-filtered in SleepSign using a 0.5–30 Hz FIR band-pass filter, and Fast Fourier Transforms (FFT; 1024 points, 4.1 s epochs, Hanning window) were applied, yielding a frequency resolution of ∼0.25 Hz. Only power values within 0.5–30 Hz were analysed. For each 4-second epoch, relative power was computed by normalizing each frequency bin to total epoch power, then averaged across all epochs of the same state (Wake, NREM, REM). Epochs containing previously detected IIS events were excluded. Data from left and right hippocampi, referenced to the same electrode, were averaged. Frontal electrode was excluded from spectral analyses due to noisy data. Spectral power was compared between genotypes for each vigilance state and channel using independent t-tests per frequency bin, with multiple comparisons corrected via False Discovery Rate (FDR).

### Homeostatic Regulation of Delta Power (Process S Simulation)

To investigate the homeostatic regulation of delta power (1-4 Hz), Process S dynamics were simulated using a well-established model^55,63,105,106^. In this model, Process S increases during Wake and REM according to a saturating exponential function and decays during NREM following an exponential decay function.

During Wake/REM:

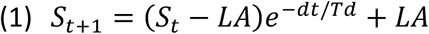

During NREM:

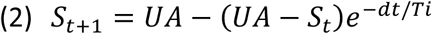

St is the simulated Process S at time t, dt the time step between consecutive measures (4 seconds), Ti and Td are time constants for increase and decay and UA and LA are the upper and lower asymptotes. The initial value S_t=0_ was also treated as a free parameter. Model parameters (Ti, Td, S_t=0_) were optimized per animal by minimizing the mean squared error between simulated and smoothed empirical delta power time series. To minimize inter-individual variability, UA and LA were fixed to 250 and 50.

Empirical delta power was calculated by collapsing power from 1–4 Hz frequency bins during NREM epochs. Epochs with power beyond ±2.5 z-score were excluded. Values were normalized within each animal as a percentage of mean NREM delta power across the 24h recording. The temporal profile was then smoothed using a centred rolling average with 1h window, requiring at least 50% valid data points per window.

### Circadian Modulation of Sigma Frequency Band

EEG power was summed across the sigma band (9–16 Hz) for each channel and vigilance state using FFT data, including only NREM epochs. Data were resampled into 1-hour bins and normalized within each animal by dividing each time point by the mean sigma power over 24 hours, centring values around 1. Sigma power was modelled for each animal and channel using a cosine function with a fixed 24-hour period:

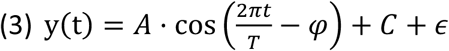

where *A* and *φ* represent the amplitude and phase (i.e. time of maximal sigma power), *C* the baseline level and *ϵ* residual noise. Model fitting was performed using the curve_fit function from scipy.optimize, with non-linear least squares optimization. Amplitudes were constrained to non-negative values and phases to the interval 0-2π. Goodness of fit was assessed using the coefficient of determination (R²).

Amplitude and R² values were averaged across channels per animal and compared using independent t-test. Phases were also averaged per animal and compared using circular statistics. For each genotype, the circular mean (μ), resultant vector length (r), and concentration parameter (κ) were computed to quantify the mean phase, phase clustering, and dispersion, respectively. Homogeneity of concentration parameters (κ) between genotypes was first tested using the equal kappa test. Differences in mean phase angles were then evaluated using the Watson–Williams test for circular means or angular randomization test (1,000 permutations) depending on the results of the equal kappa test.

### Slow-Waves Detection

Slow-waves (SWs) during NREM sleep were automatically detected using the sw_detect() function from the YASA Python package^107^. Prior to SWs detection interictal spikes IIS were removed. SWs were defined by the following criteria: negative peak duration 125 ms–1 s, amplitude 1–3 z-units; positive peak duration 125 ms–1 s, amplitude 0.5–3 z-units; and peak-to-peak amplitude 2.5–6 z-units. For each animal and channel, SW counts were normalized to total NREM time to compute SW density. Peak-to-peak amplitude, duration, and slope were extracted for each animal and compared between genotypes using t-tests, Welch’s t-tests, or Mann–Whitney tests as appropriate.

### Spindle Detection

Spindle detection during NREM sleep was performed using the spindles_detect() function from the YASA Python package, based on the A7 algorithm^108^. The detection combined three criteria: 1) the ratio of sigma (9–16 Hz) to broadband (1–30 Hz) power, thresholded at the 75th percentile; 2) the Pearson correlation (r ≥ 0.65) between sigma-and broadband-filtered signals and 3) the moving root mean square of the sigma-filtered signal, thresholded at 2.5 SD above the mean.

Only events lasting 0.4–3 s were retained^109^, and spindles occurring within 500 ms were merged. For one WT and two HET animals no spindles were detected, these were excluded from all subsequent analyses. For each animal, the density (spindles per minute), mean amplitude, mean duration and mean number of oscillations were calculated and compared between genotypes using t-tests, Welch’s t-tests, or Mann–Whitney tests as appropriate. Spindle coherence between channels was estimated as the proportion of overlapping events between channels (≤0.5 s window) using compare_channels().

### Infra-Slow Oscillation Analysis of Sigma Power

Infra-slow fluctuations in sigma power during uninterrupted long NREM episodes (≥96s) were analysed as described previously^58,110^. For each channel, sigma power was estimated using a Morlet wavelet transform (tfr_array_morlet, MNE) with 0.2Hz frequency resolution and 4-cycle wavelets. Power was collapsed across frequencies per timepoint, mean-centred, and resampled at 10Hz. For visualization, the 10Hz resampled sigma signal was low-pass filtered (FIR, order 100, cutoff 0.025 Hz).

The resulting sigma signal was transformed via FFT to obtain the infra-slow power spectrum (0–0.12 Hz). Each spectrum was normalized by (1) episode length and (2) mean power across 0–0.12 Hz frequencies. Because different episode lengths result in different frequency resolution, the normalized spectra were binned into 0.005 Hz-wide bins and averaged across episodes. Spectra were further normalized by subtracting mean power in the baseline plateau (0.08–0.12 Hz). The relative power within the 0.01–0.04 Hz band was collapsed and compared with an independent t-test.

### Microarousal Analysis and Phase Coupling to Infra-Slow Oscillations

Long NREM episodes (≥96s) containing brief interruptions (≤16 s and <30% of total episode duration) were detected, and infra-slow fluctuations in sigma power were analysed as described above. The instantaneous phase of the low-pass filtered sigma signal (FIR, order 100, cutoff 0.025 Hz) was extracted using the Hilbert transform^61^. Phases were circularly aligned such that sigma peaks corresponded to 0° and troughs to 180°. The phase of each microarousal (MA) onset within NREM episodes was interpolated from the sigma phase time series. For each animal, mean preferred phase (μ) and phase concentration (r) were computed using circular statistics, and phase distributions were tested for non-uniformity with the Rayleigh test. MA density was calculated as the number of MAs per hour of NREM sleep and compared between genotypes using independent t-tests.

### Tissue Fractionation and Purification of Synaptic Fractions

Preparation of synaptic fractions followed a previously described protocol^62^. Tissue was homogenized (9:1 v/w) using 1 mL glass-glass tissue grinders (Borosilicate Dounce homogenizer, Wheaton) with 20 strokes on ice in homogenization buffer (HB: 50 mM Tris, 0.32 M Sucrose, 5 mM EDTA, 1 mM EGTA, 20 uM ZnCl2, 50 mM NaF, 1 mM Na-Orthovanadate, 2.5 mM Na-Pyrophosphate, 1 ug/ml Aprotinin, 1ug/ml Leupeptine, 1/2500 PMSF, pH 7.4). The homogenate was centrifuged at 1,400×g for 10 minutes at 4°C using an Eppendorf 5417R centrifuge. The supernatant was retained, and the pellet was re-homogenized twice under the same conditions. 150 μL of the first supernatant were aliquoted to use in MS experiments as a nuclei-depleted homogenate. The combined supernatants from all three rounds of homogenization were centrifuged at 700×g for 10 minutes. The resulting supernatant was centrifuged at 21,000×g for 30 minutes at 4°C and the pellet resuspended in HB and layered onto a 0.85/1/1.2 M sucrose gradient in a 11×60 mm tube (Beckman Coulter, CA, USA). The gradient was centrifuged at 82,500×g for 2 hours at 4°C using a Sw60 Ti rotor (Beckman Coulter) in an ultracentrifuge (Optima L-90K Ultracentrifuge, Beckman Coulter). Synaptosomes were collected from 1-1.2M interface, diluted to 10% sucrose with 50 mM Tris buffer (pH 7.4) and centrifuged at 21,000×g for 30 minutes at 4°C using an Eppendorf 5117R rotor. The resulting pellet was resuspended in 50 mM Tris buffer (pH 7.4) containing 0.5% Triton X-100, incubated on ice for 15 minutes and centrifuged again (21,000×g, 30 min, 4°C). The Triton X-100 insoluble fraction was used for MS experiments.

Homogenate and synaptic fractions were extracted in 5% SDS and 50 mM Tris-HCl (pH 8) for 10 minutes at 37°C with gentle shaking, then centrifuged at 10,000×g for 1 minute, and supernatants was collected. Protein was quantified using a micro-BCA assay (Thermo Fisher Scientific, Waltham, MA, USA) following the standard microplate procedure. Absorbance measurements were taken at 560 nm using the xMark™ Microplate Absorbance Spectrophotometer (Bio-Rad).

### Sample Preparation for MS

Mass spectrometry (MS) sample preparation was carried out using a modified version of the DNA micro spin column suspension trapping protocol described by Thanou et al., 2023^111^. 20 µg of protein was combined with S-Trap lysis buffer (5% SDS, 50 mM Tris-HCl, pH 8.5) supplemented with 5 mM tris(2-carboxyethyl)phosphine (TCEP) and 20 mM chloroacetamide (CAA). Reduction and alkylation were performed sequentially, with samples incubated at 95°C for 15 minutes followed by 55°C for 15 minutes in a thermomixer set at 1500 RPM.

After treatment, samples were acidified to a final phosphoric acid concentration of 1.1% (from a 12% stock solution) and mixed with six volumes of binding/washing buffer (90% methanol, 100 mM Tris-HCl, pH 8). The mixture was then loaded onto a plasmid DNA micro column (HiPure, Magen Biotechnology). Proteins were retained on the column by centrifugation at 1400×g for 1 minute, followed by four washes with binding/washing buffer.

Columns were subsequently transferred to new LoBind tubes (Eppendorf), where digestion was carried out by adding 0.8 µg of Trypsin/Lys-C (Promega) in 50 mM NH₄HCO₃. Samples were incubated overnight at 37°C in a humidified incubator. Tryptic peptides were sequentially eluted using 50 mM NH₄HCO₃, 0.1% formic acid, and 0.1% formic acid in acetonitrile. The collected peptides were dried using a SpeedVac and stored at −80°C until further analysis.

### LC-MS Analysis

Each tryptic digested sample was reconstituted in 0.1% formic acid, and peptide concentration was measured using a tryptophan fluorescence assay^112^. A total of 150 ng of peptides was loaded onto an Evotip Pure (Evosep) before chromatographic separation. Peptide separation was performed using the Evosep One liquid chromatography system following the standardized 30 samples per day method. The separation was carried out on a 15 cm × 150 μm reverse-phase column packed with 1.5 µm C18 beads (EV1137, Evosep), which was connected to a 20 µm ID ZDV emitter (Bruker Daltonics).

Peptides were introduced into the timsTOF HT mass spectrometer (Bruker Daltonics) via electrospray ionization using a CaptiveSpray source. The mass spectrometer was operated with the following settings: scan range of 100–1700 m/z, ion mobility range of 0.65–1.5 Vs/cm², ramp time of 100 ms, and accumulation time of 100 ms. Collision energy was dynamically adjusted, decreasing linearly with inverse ion mobility from 59 eV at 1.6 Vs/cm² to 20 eV at 0.6 Vs/cm².

Data were acquired in dia-PASEF mode, with each acquisition cycle lasting 1.38 seconds and consisting of one MS1 full scan followed by 12 dia-PASEF scans. Each dia-PASEF scan included two isolation windows, collectively covering an m/z range of 300–1200 and an ion mobility range of 0.65–1.50 Vs/cm². The placement of dia-PASEF isolation windows was optimized using the py-diAID tool^113^. Ion mobility calibration was automatically performed at the start of each sample using the following calibrant ions (m/z, 1/K₀): 622.029, 0.992 Vs/cm²; 922.010, 1.199 Vs/cm²; and 1221.991, 1.393 Vs/cm².

### Peptide Identification and Protein Mapping

DIA-PASEF raw data were analysed using DIA-NN 1.8.1^114,115^. A predicted spectral library was generated in silico from the UniProt mouse reference proteome (SwissProt and TrEMBL, including canonical and additional isoforms, release 2023-5). Peptide digestion was simulated using Trypsin/P, allowing up to one missed cleavage. Carbamidomethylation was set as a fixed modification, while oxidation and N-terminal methionine excision (with a maximum of one modification per peptide) were specified as variable modifications. Peptide length was restricted to 7–30 amino acids, precursor charge state was set between 2–4, and precursor m/z was limited to 280–1220. Both MS1 and MS2 mass accuracy were set at 15 ppm, with a scan window of 9. The double-pass mode and match-between-runs feature were enabled, with protein isoforms used for protein inference. All other parameters remained at default settings.

For downstream analysis, MS-DAP 1.0.8^116^ was used to process the DIA-NN output. Peptide-level filtering was set to retain only peptides that were confidently detected in at least 75% of samples per group. Peptide abundance values were normalized using the VSN (Variance Stabilizing Normalization) algorithm, followed by mode-between normalization at the protein level. One outlier sample was identified in the quality control analysis generated by the MS-DAP report in the synaptic fractions dataset and two in the Homogenate dataset. These outliers were excluded from further analyses.

Additionally, for each protein, values within a specific ZT group and genotype were cleaned using the Interquartile Range (IQR) method. The IQR was calculated as the difference between the 1st (Q1) and 3rd (Q3) quartile of the data and any values falling below Q1 – 1.5·IQR or above Q3 + 1.5·IQR were removed.

### Model Selection Analysis for Temporal Regulation Identification

To investigate rhythmic protein expression across genotypes over time we implemented a model selection framework^63^ with 7 different models:

- Null Model: Assumes constant expression over time (no rhythmicity).

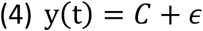
- Shared Cosine Model: Captures oscillatory patterns with shared amplitude and phase between genotypes. Follows eq. 3.
- WT-Specific Model: Assumes oscillatory pattern in WT but constant expression in HET.

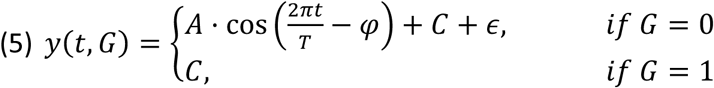
- HET-Specific Model: Assumes constant expression in WT but oscillatory pattern in HET.

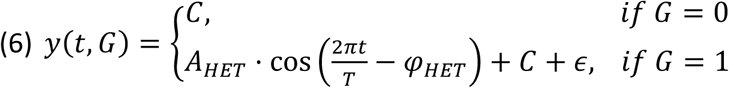
- Genotype-Dependent Amplitude Model: Fits different amplitudes for WT and HET while sharing a common phase.

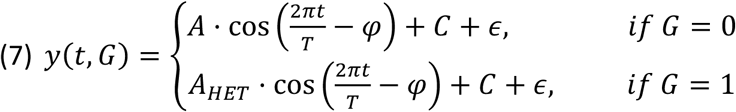
- Genotype-Dependent Phase Model: Fits different phases for WT and HET while keeping amplitude constant.

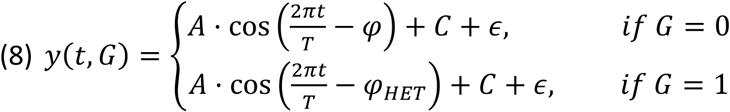
- Fully Genotype-Dependent Model: Allows both amplitude and phase to vary by genotype.

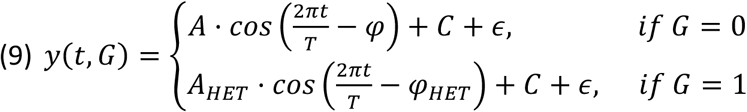

In all cases, the period (T) was fixed to 24 h, *C* represents the baseline expression level, and ɛ residual noise. *A* and *φ* represent the amplitude and phase (i.e. time of maximal abundance) derived from WT fitting. In some equations, these parameters are shared with HET, while in others the parameters *A_HET_* and *φ_HET_* were introduced to account for genotype differences. In those cases, the genotype variable (G) was encoded as G=0 for WT and G=1 for HET mice.

Prior to model fitting, protein expression values (log₂-transformed) were normalized by subtracting the genotype specific mean to remove baseline differences. Model fitting was implemented using non-linear least squares optimization (curve_fit, scipy.optimize). Amplitudes were constrained to non-negative values and phase parameters restricted to 0-2π. Model fits assumed independent Gaussian errors with constant variance. For each protein, the sum of squared residuals was used to compute the Bayesian Information Criterion (BIC; Astropy function bayesian_info_criterion_lsq). Schwarz weights were derived from BIC values to quantify model support, with the highest-weight model defining the best fit^63^.

To refine detection of genotype-specific rhythms, proteins initially classified as non-rhythmic (Null model) but showing a WT- or HET-specific cosine model as the second-best fit were reanalysed using only data from the corresponding genotype. BIC-based Schwarz weights were recalculated, and proteins were classified as rhythmic in WT or HET when the genotype-specific cosine outperformed the Null model.

### Decile-Based Enrichment Analysis of Subthreshold Trends

To determine whether genotype-specific proteins reflect real biological changes instead of merely subthreshold rhythms in the other genotype, cosine models were independently fitted to expression profiles from WT and HET animals using constrained non-linear least squares (eq. 3). For each genotype, the R² was computed as a quantitative measure of goodness-of-fit and used to rank proteins (excluding genotype-specific rhythmic proteins) and divide into deciles. The distribution of WT- and HET-specific rhythmic proteins across R² deciles of the opposite genotype was then assessed using a hypergeometric test to evaluate enrichment in particular deciles. This approach allowed us to test whether proteins rhythmic in one genotype tend to exhibit some degree of oscillatory behaviour in the other genotype but fail to reach statistical significance due to increased noise or variability. As a positive control, proteins rhythmic in both genotypes (shared, phase-shifted, or amplitude-differential) were analysed similarly.

### Pathways and GO Terms Enrichment Analysis

To explore the biological relevance of the rhythmic protein subsets identified by model selection, Pathway and Gene Ontology (GO) enrichment analyses were performed using the pathfindR package in R^117^, following a previously published approach^62^.

PathfindR first builds a Protein Interacting Network (PIN) from all differentially expressed (DE) molecules (genes/proteins), or oscillatory proteins in this case, using a protein–protein interaction (PPI). A high-confidence PPI network was constructed by integrating data from two complementary sources: STRING v12.0 and BioGRID v4.4.237. STRING interactions for *Mus musculus* and *Homo sapiens* with combined confidence ≥0.9 were retrieved, while BioGRID physical interactions were filtered to retain only same-species pairs. Protein identifiers were mapped to gene symbols using biomaRt^118,119^, redundant and low-quality entries (self-interactions, reciprocal duplicates) were removed, and homologous mappings between species were standardized. The final integrated PPI network contained 972,209 unique interactions.

Active subnetworks were then identified using a greedy search algorithm constrained to a maximum depth of 1, thereby including only direct neighbours of oscillatory proteins. Subnetworks with ≥10 molecules were retained and overlapping ones (≥50% shared genes) were filtered to keep the highest-scoring network. Pathway enrichment was then performed for each retained subnetwork using a one-sided hypergeometric test, with the full PIN used as background.

Pathways investigated with pathfindR were taken from MSigDB (https://www.gsea-msigdb.org). From the C2 collection, only REACTOME, WikiPathways, and KEGG terms were used (3,224 gene sets); from the C5 collection, all GO Biological Process (BP), Cellular Component (CC), and Molecular Function (MF) terms were included (10,454 sets). As the greedy search is stochastic, the analysis was repeated 50 times, retaining only terms enriched in ≥13 iterations (>25% occurrence). Additionally, terms containing <3 proteins were discarded. To avoid biologically irrelevant enrichments in our context, terms related to viral infections (HIV, SARS-CoV-2) were also excluded. Finally, to reduce complexity and facilitate interpretation, enriched terms were clustered using pathfindR’s cluster_enriched_terms() function, and only the most representative term from each cluster was reported. This was done separately for Pathways and for each GO sub-ontologies (BP, CC, MF).

### Temporal Coherence Filtering of Pathway and GO Terms

To incorporate temporal synchronization of proteins within enriched Pathways and GO Terms, the circular standard deviation (Std) of protein peaks within each term were analysed. A threshold of 3.13 circular Std was applied to identify pathways in which ≥80% of proteins oscillated within an 8-hour window centred around the mean (±4 h), assuming a normal distribution. This cutoff corresponds to twice the 4-hour sampling interval, ensuring detection of pathways with tightly synchronized oscillations while accounting for biological and technical variability.

### Sleep Deprivation

Sleep deprivation (SD) was achieved by novel environment and novel object exposure. Animals were transfer to a novel cage containing unfamiliar objects at ZT0. The procedure was continuously monitored by an experimenter to ensure the maintenance of wakefulness throughout the deprivation period. If behavioural signs of sleep were observed, additional novel objects were introduced to re-engage exploratory activity. If animals continued to display sleep-like behaviour, mild sensory stimulation was applied, including gentle tapping on the cage or disturbance of the bedding material. At ZT4 animals were sacrificed and cortices were collected for further analysis.

### Immunoblot analysis

After isolation of synaptic fractions from cortical tissue, protein concentrations were quantified using a micro bicinchoninic acid (BCA)-based protein assay kit (Thermo Fisher Scientific, Waltham, MA, USA), mixed with 10xLSB (500 mM Tris, 20% (w/v) SDS, 10% (v/v) beta-mercaptoethanol, 0.1% (w/v) bromophenol blue, pH 7.4) and boiled at 95°C for 5 minutes. The prepared samples were then boiled at 95°C for 5 minutes. Aliquots were subsequently generated and stored at –80°C until further use. Proteins were separated by electrophoresis using Fast-Casting TGX Stain-Free gels (Bio-Rad) in a vertical Mini-PROTEAN system kit (Bio-Rad). Following electrophoresis, the polyacrylamide gels were activated under UV light for 2 minutes using a ChemiDoc Imaging System (Bio-Rad), Next, transfer was conducted in a vertical Mini-PROTEAN system kit (Bio-Rad), using a custom-made transfer buffer (48 mM Tris, 39 mM Glycine, 0.04% (w/v) SDS, 20% (v/v) Methanol, pH 9), at a constant voltage of 100 V for 90 minutes. PVDF membranes were then fixed following in pure acetone at 4°C for 30 minutes with continuous agitation and dried for 30 minutes at 60°C^120^.

Following fixation, membranes were reactivated in methanol and washed in TBS-T buffer (50 mM Tris, 150 mM NaCl, 1% (v/v) Tween-20, pH 7.4) for 5 minutes. The membranes were then blocked for 15 minutes using a commercial solution (EveryBlot Blocking Buffer, Bio-Rad, #12010020) and incubated with the primary antibody (1:500 goat anti-Baiap2 (ab15697 Abcam); 1:1000 mouse anti-Grin2b (610417 BD Bioscience); 1:1000 rabbit anti-Dlg3 18036-1-AP ProteinTech)) diluted in TBS-T for either 2 hours at room temperature (RT) or overnight at 4°C. Following incubation, membranes were washed three times for 5 minutes each with TBS-T and then incubated for 1 hour at RT with a secondary antibody conjugated to an infrared fluorophore (1:7500 IRDye® 680RD anti-Rabbit (9226-68073 Licor-Bio), IRDye® 800CW anti-Mouse (9226-32212 Licor-Bio) or IRDye® 800CW anti-Goat (9226-32214 Licor-Bio)). Membranes were next washed three more times for 5 minutes each with TBS-T, followed by a final wash in TBS. Membranes were imaged using an LI-COR 9120 Odyssey or Odyssey M Infrared Imaging System for infrared fluorescence detection. Band intensity was analysed using Image Studio Lite v5.2 (Li-Cor), and total protein quantification previously obtained was used for normalization.

### Real Time – quantitative Polymerase Chain Reaction (RT-qPCR)

For RNA experiments, nuclear fractions obtained during the brain fractionation protocol were used. RNA was extracted using the RNeasy Lipid Tissue Mini Kit (Qiagen) following the manufacturer’s recommended protocol. The RNA integrity number (RIN) was measured using an Agilent 2100 Bioanalyzer, quantified using a Nanodrop 2000 (ND-200, Thermo Scientific) and transcribed into cDNA. To quantify gene expression, a total of 100 ng of cDNA was used for each reaction on a 384-well plate (Thermo Fisher, AB1384). The reaction mixture included 2 μL of 50 ng/ μL of cDNA, 5 μL of PowerUp SYBR Green Master Mix (Applied Biosystems™), 300 nM of specific primers (see **Table 2**), and DNA- and RNA-free H₂O, making up a final volume of 10 μL. The plate was then analyzed using a 7900HT Fast Real-Time PCR System with 384-Well Block Module (Applied Biosystems™) in combination with the software Sequence Detection System v2.4 Enterprise Edition (SDS 2.4; Applied Biosystems™). All samples were analyzed in triplicate for the gene of interest and the reference gene *Gapdh*, which was used to normalize values gene expression following the equations:

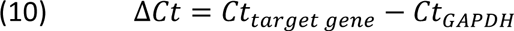

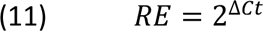

For each sample, the Mean and Standard Deviation (Std) of the Ct values within technical replicates were calculated. Whenever the Std exceeded 0.4, Z-scores were calculated, and the replicate with the highest value was removed to eliminate outliers resulting from technical errors.

## Data Availability

All the data generated by the bioinformatics analysis performed in this manuscript can be found in the supplementary tables. Mass spectrometry proteomics data has been deposited to the ProteomeXchange Consortium via the PRIDE partner repository121 with the dataset identifier PXD073485.

## Code Availability

All custom-made code is available from GitHub:

https://github.com/Molecular-Physiology-of-the-Synapse/synapse_sleep_2026/releases/tag/v1.0

## Author Contributions

DdCB, OZR, AP, CVR, ACG, DAA, GE, DZ, TV, CR, FK, RVK and BDV performed experiments. DdCB, MAD, JS, GR, ABS and AB designed and supervised all studies. MAD, JS, GR, ABS and AB secured funding. DdCB and AB wrote the manuscript. All authors reviewed and approved the manuscript.

## Acknowledgments

DdCB, OZR, AP, ACG, DAA, GE and AB financial support was provided by: PID2024-160538OB-I00 and PID2021-124411OB-I00 (MICIU/AEI/10.13039/501100011033/ and FEDER-EU), Award AC17/00005 by ISCIII through AES2017 and within the NEURON framework, Ramón y Cajal Fellowship (RYC-2011-08391p), IEDI-2017-00822, Universitat Autònoma de Barcelona and AGAUR (2017 SGR 1776 and 2021 SGR 01005). BDV and MAD for the financial support provided by: MCIU/AEI/FEDER Grants PID2024-162613OB-I00; PID2021-127044OB-I00. DdCB and ACG thank AGAUR/Generalitat de Catalunya/FEDER, EU for ‘Ajuts per a la contractació de personal investigador novel (FI)’ Ref. 2020FI_B00130 and 2022 FI_B 00996. DdCB thanks Fundació Universitària Agustí Pedro i Pons for ‘Ajuts per a estudis o projectes fora de Catalunya’. OZR thanks the Universitat Autonoma de Barcelona for Investigador Predoctoral en formación (PIF-UAB) Ref. B21P0024. DdCB, OZR, AP, ACG, DAA, GE and AB thank the CERCA Programme/Generalitat de Catalunya for institutional support. The study was supported by the generous funding provided by the association of Spanish SYNGAP1-DEE families, SYNGAP1 España.

---

**Figure S1.**
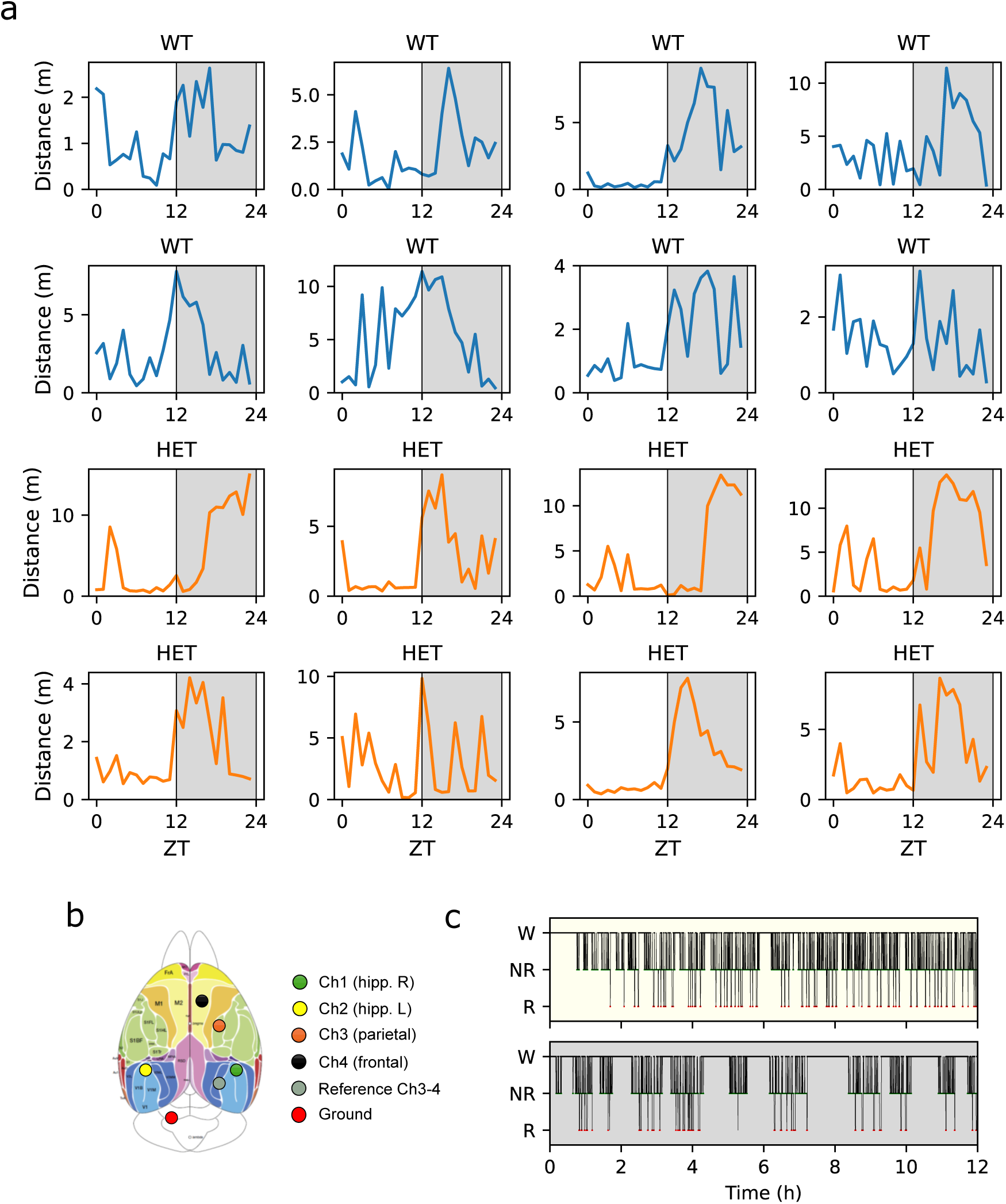
Video-tracking of locomotor activity, EEG experimental set-up and temporal distribution of sleep/wake states. **(a)** Activity patterns of WT (blue; top two rows) and HET (orange; bottom two rows) mice over a 24-hour period as measured by the distance travelled (m) per hour. Grey shaded areas indicate the dark period of the light/dark cycle. **(b)** Electrode implantation scheme showing the stereotaxic placement of 8 electrodes resulting in 4 EEG channels: frontal cortex (black), parietal cortex (orange), and bilateral hippocampus (green and yellow). Adapted from: Kirkcaldie, 2012^123^. **(c)** Representative hypnogram from a wild-type (WT) mouse displaying manual scoring of vigilance states (Wake, NREM, REM) across the light (top) and dark period (bottom).

**Figure S2.**
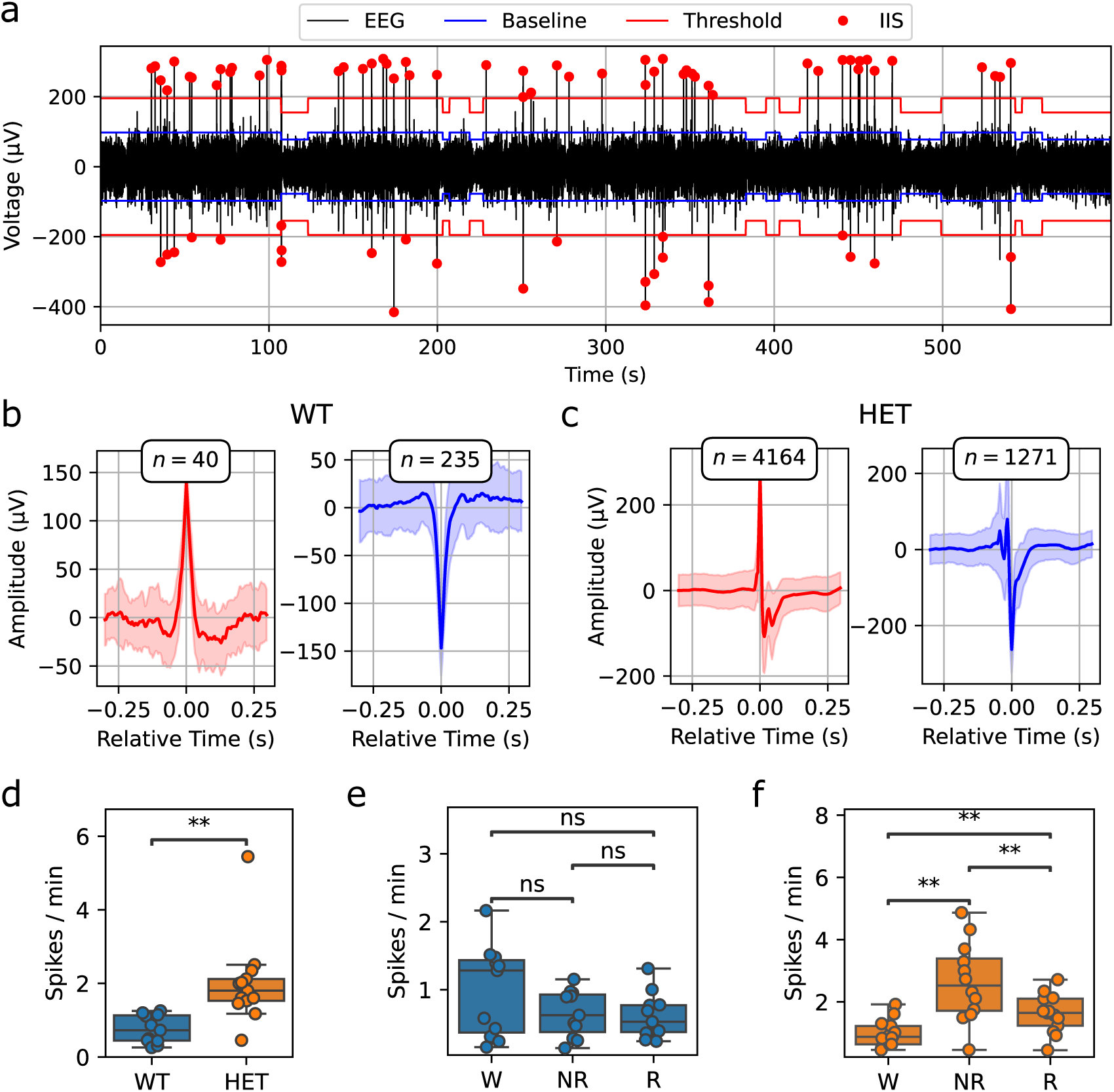
Detection and analysis of Interictal Spikes (IIS). **(a)** Example of IIS detection performance in a 10-minute period with high IIS activity on a HET mouse. The blue line represents the 99th percentile of the absolute amplitude of the signal within each behavioural state (baseline), while the red line represents the threshold for spike detection (2·baseline). Detected spikes are marked by red dots. **(b, c)** Average waveforms of positive (left) and negative (right) spikes detected in a WT (b) and HET (c) animal for a given channel. Spikes are aligned to their peak (time 0). Shaded areas represent the standard deviation. The total number of detected events for each category is indicated above each plot. **(d)** Comparison of IIS frequency between genotypes. Data are presented as boxplots with individual data points overlaid. Boxplots show median, interquartile range (IQR), and 1.5×IQR whiskers. Statistics: independent t-test, WT n=11, HET n=13, p=0.001. **(e, f)** Comparison of IIS frequency between behavioural states in WT (e) and HET (f). Statistics: paired t-test or Wilcoxon signed-rank test depending on data normality, followed by Holm correction. WT n=11, HET n=13. WT: non-significant. HET: W vs NR adj. p-value = 0.003, NR vs R adj. p-value = 0.003, W vs R adj. p-value = 0.006. Significant differences are indicated as follows: *p < 0.05, **p < 0.01, ***p < 0.001.

**Figure S3.**
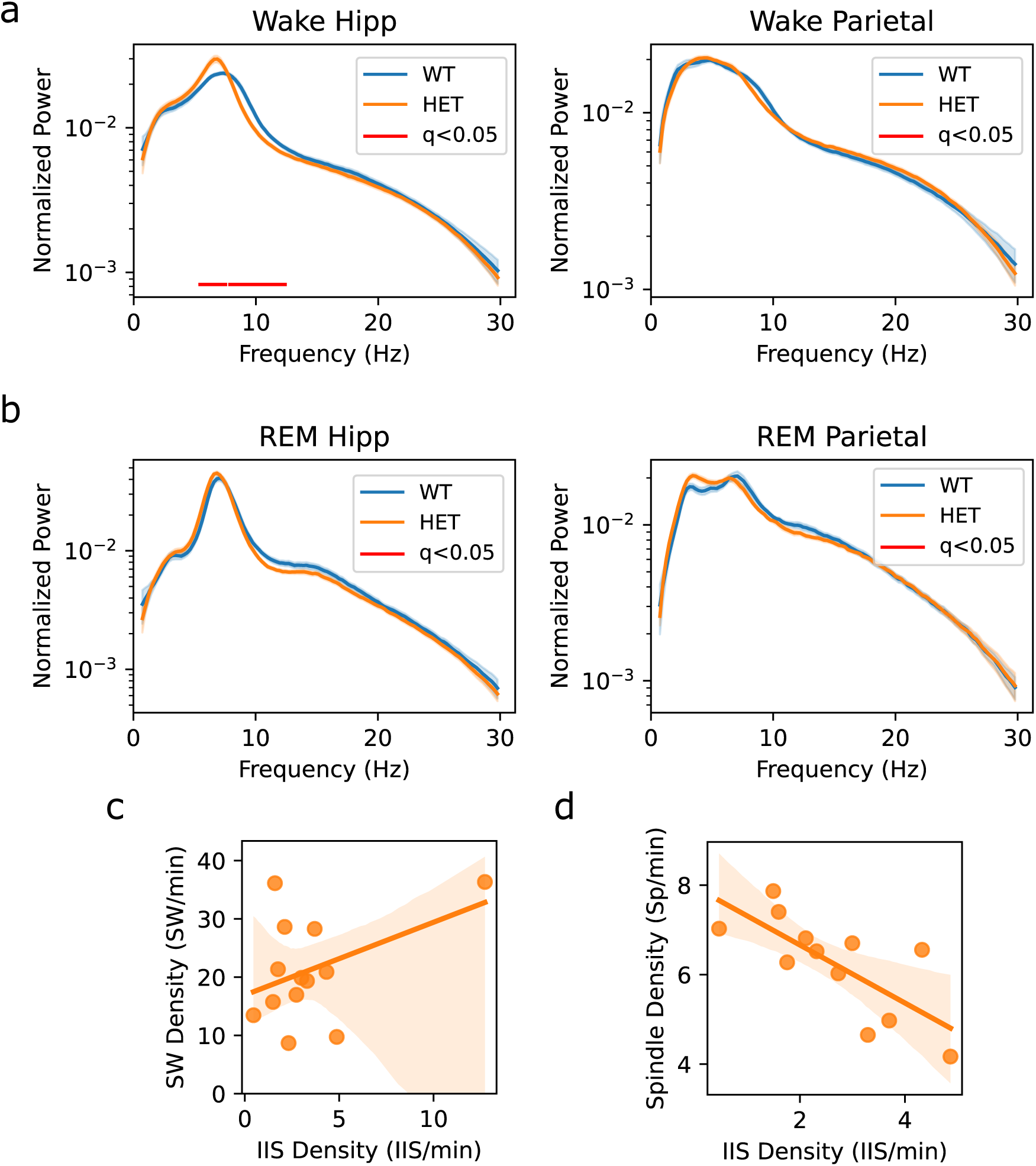
Power spectra analysis of wake and REM sleep and correlations between NREM oscillations and IIS. **(a, b)** Normalized power spectra during Wake (a) and REM (b) after excluding IIS epochs for both WT (blue) and HET (orange) animals in hippocampal channels (left panel) and parietal channel (right panel). The shaded areas around the line plots represent the SEM. Red lines below the traces indicate frequencies with significant differences (multiple independent t-test with Benjamini-Hochberg (BH) FDR correction, q < 0.05). **(c)** Correlation between SW density and IIS density in HET mice. Pearson r = 0.43, p = 0.14. **(d)** Correlation between spindle density and IIS density in HET mice. Pearson r = - 0.74, p = 0.006.

**Figure S4.**
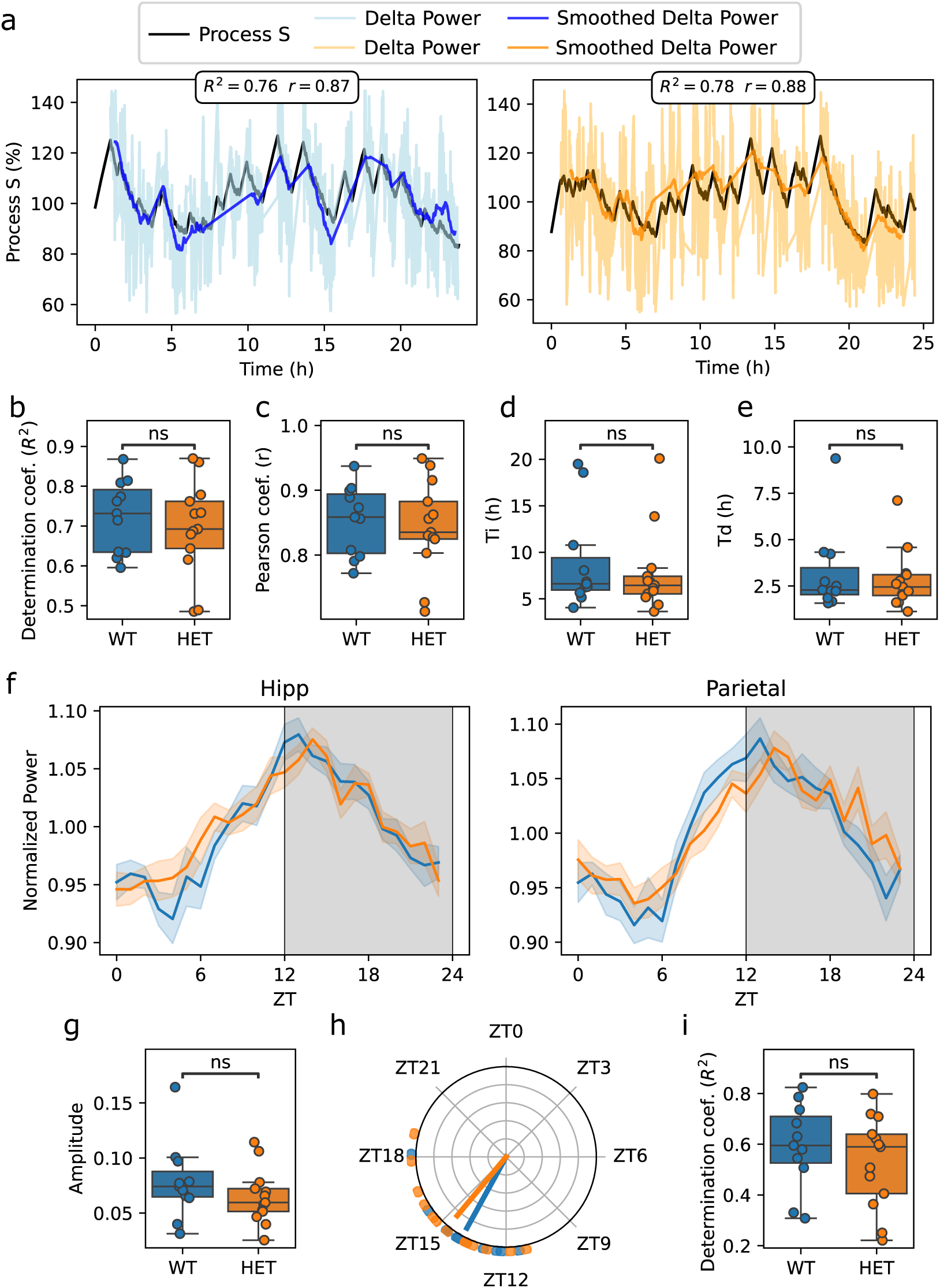
Homeostatic regulation of delta and circadian regulation of sigma bands. **(a)** Representative examples of Process S simulation and comparison with empirical delta power in a WT (left, blue) and a HET (right, orange) animal. Process S (black line) was simulated using established methods with fixed asymptotes (UA = 250, LA = 50). Dark blue (WT) and orange (HET) traces represent the smoothed delta power using a 1h rolling window. Lighter traces correspond to delta power calculated with a 2 min rolling window. The R^2^ and Pearson’s r for each animal are indicated. **(b, c)** Comparison of model fit across genotypes using R^2^ (b) and Pearson r (c) calculated for each animal. Statistics: independent t-tests: WT n=11, HET n=13, *R^2^* p=0.49, *r* p=0.68. **(d, e)** Comparison of optimized time constants Ti (d) and Td (e) between WT and HET mice. Values are shown as boxplots with individual data points overlaid. Statistics: Mann-Whitney U tests: WT n=11, HET n=13, Ti p=0.73, Td p=0.86. **(f)** Average 24-hour dynamics of sigma power during NREM sleep for WT (blue) and HET (orange) mice in the hippocampal (left) and parietal (right) channels. Data are plotted as mean ± SEM for each genotype. Grey shaded area represents the dark period of the light/dark cycle. **(g)** Comparison of amplitude of sigma oscillations between genotypes. Statistics: independent t-test, WT n=11, HET n=13, p=0.27. **(h)** Comparison of the time of peak of the 24-h sigma power rhythm. Dots outside the circle represent the phase of individual animals obtained (blue = WT, orange = HET). Arrows indicate the genotype mean phase, with arrow length proportional to the mean resultant length (r, i.e. phase concentration). Statistics: Watson-Williams Test, WT n=11, HET n=13, p=0.41. **(i)** Comparison between genotypes of R² values obtained from the cosine fitting. Statistics: independent t-test, WT n=11, HET n=13, p=0.39. Boxplots show median, interquartile range (IQR), and 1.5×IQR whiskers.

**Figure S5.**
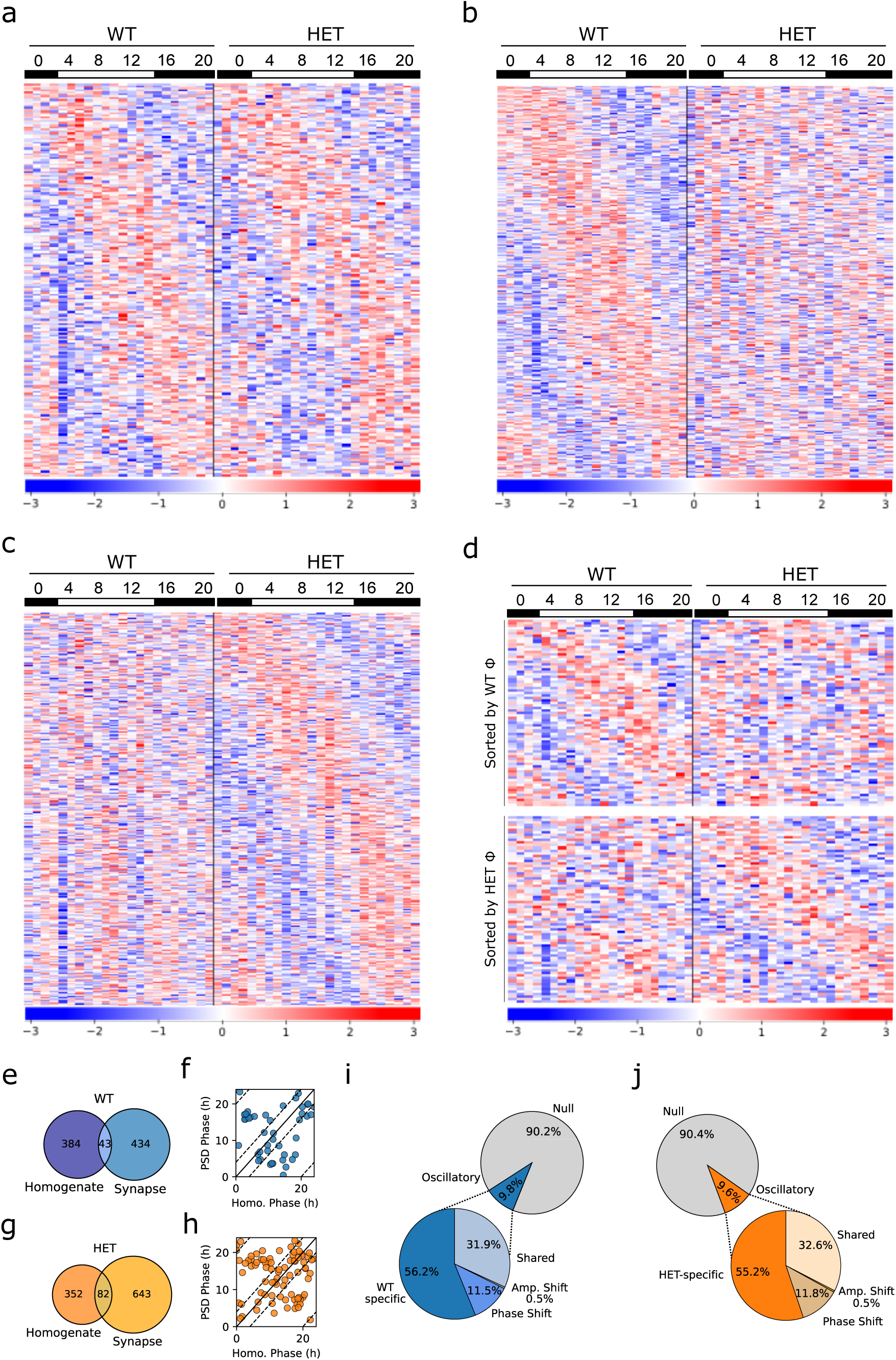
Daily expression dynamics of homogenate proteins. **(a-d)**. Heatmaps displaying normalized expression of cycling proteins from the following expression models: shared-regulation (a), WT-specific (b), *Syngap1*^+/-^-specific (c) and phase-shift (d). Z-scored protein abundances are color-coded, with red indicating higher expression and blue indicating lower expression. Proteins (rows) were sorted by their phase. **(e)** Venn diagram showing the overlap between WT cycling proteins in homogenates and synaptic fractions. **(f)** Scatter plots comparing the phase of WT cycling proteins in both compartments. Dashed lines indicate a ±4-hour window around the identity line. **(g)** Venn diagram showing the overlap between WT cycling proteins in homogenates and synaptic fractions. **(h)** Scatter plots comparing the phase of WT cycling proteins in both compartments. Dashed lines indicate a ±4-hour window around the identity line. **(i, j)** Pie chart showing the proportion of proteins best fit by each of the tested models in homogenates for WTs (i) and HETs (j). No proteins were classified in the model with simultaneous phase and amplitude shift.

**Figure S6.**
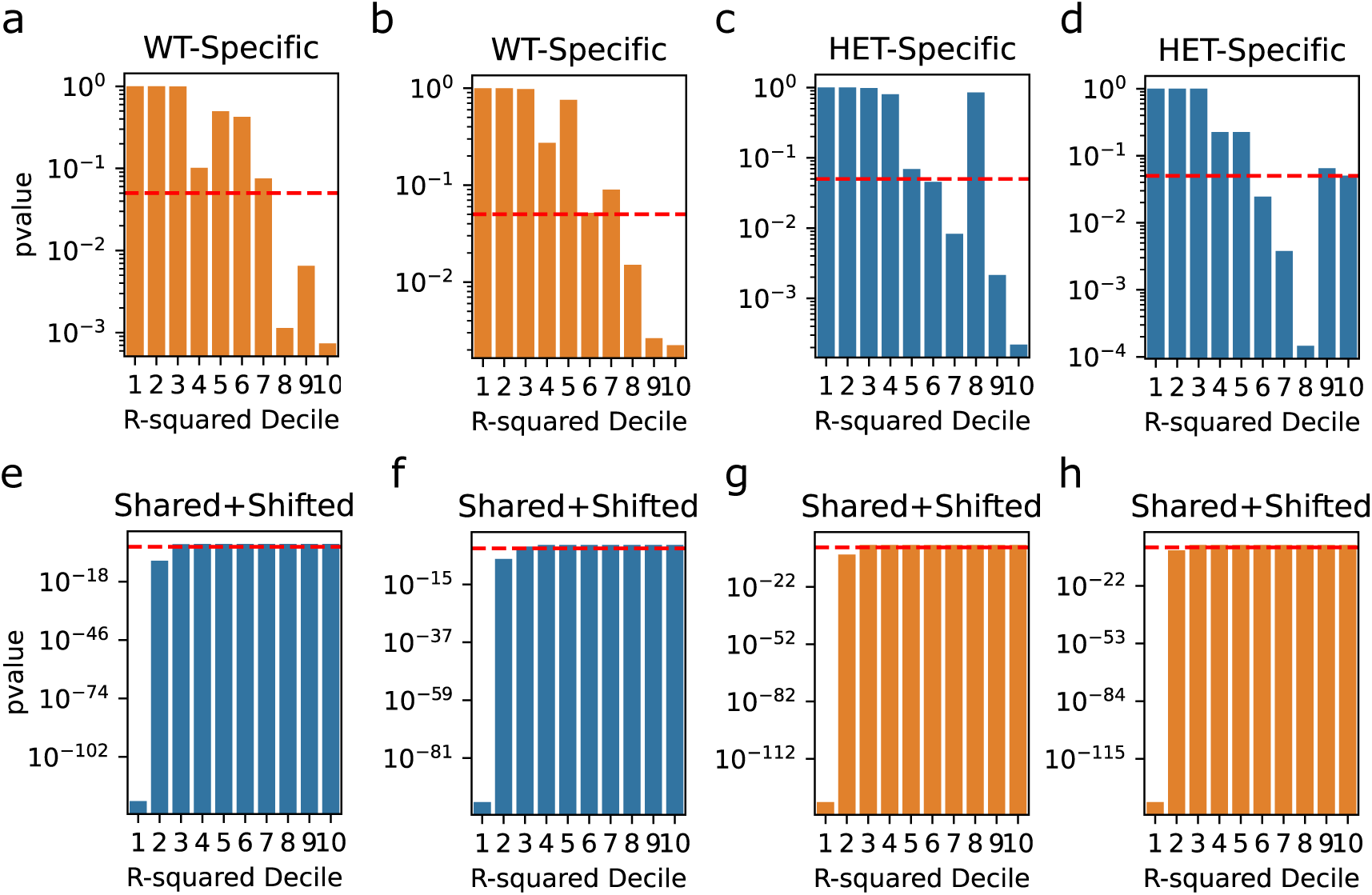
Enrichment analysis of genotype-specific rhythmic proteins across R² deciles in the opposing genotype. **(a, b)** Enrichment analysis of WT-specific cycling proteins in homogenates (a) and synapses (b) across R² deciles in HET-ranked proteins of their respective compartment. **(c, d)** Enrichment analysis of HET-specific cycling proteins in homogenates (c) and synapses (d) across R² deciles in WT-ranked proteins of their respective compartment. **(e-h)** Enrichment analysis of cycling proteins in both genotypes (Shared oscillations + Phase-shifted + Amp. Shifted) in R² deciles in WT homogenates (e), WT synapses (f), HET homogenates (g) and HET synapses (h). The red dashed line represents the significance threshold (p = 0.05). The y-axis displays the p-value of the hypergeometric test for each decile, plotted on a logarithmic scale.

